# GADMA2: more efficient and flexible demographic inference from genetic data

**DOI:** 10.1101/2022.06.14.496083

**Authors:** Ekaterina Noskova, Nikita Abramov, Stanislav Iliutkin, Anton Sidorin, Pavel Dobrynin, Vladimir Ulyantsev

**Affiliations:** Computer Technologies Laboratory, ITMO University, St. Petersburg, 197101, Russia; HSE University, St. Petersburg, 194100, Russia; Laboratory of Biochemical Genetics, St. Petersburg State University, St. Petersburg, 199034, Russia; Human Genetics Laboratory, Vavilov Institute of General Genetics RAS,Moscow, 119991, Russia

**Keywords:** demographic inference, population genetics, genetic algorithm, hyperparameter optimization

## Abstract

**Background:** Inference of complex demographic histories is a source of information about events that happened in the past of studied populations. Existing methods for demographic inference typically require input from the researcher in the form of a parameterized model. With an increased variety of methods and tools, each with its own interface, the model specification becomes tedious and error-prone. Moreover, optimization algorithms used to find model parameters sometimes turn out to be inefficient, for instance, by being not properly tuned or highly dependent on a user-provided initialization. The open-source software GADMA addresses these problems, providing automatic demographic inference. It proposes a common interface for several likelihood engines and provides global parameters optimization based on a genetic algorithm.

**Results:** Here, we introduce the new GADMA2 software and provide a detailed description of the added and expanded features. It has a renovated core code base, new likelihood engines, an updated optimization algorithm and a flexible setup for automatic model construction. We provide a full overview of GADMA2 enhancements, compare the performance of supported likelihood engines on simulated data and demonstrate an example of GADMA2 usage on two empirical datasets.

**Conclusions:** We demonstrate the better performance of a genetic algorithm in GADMA2 by comparing it to the initial version and other existing optimization approaches. Our experiments on simulated data indicate that GADMA2’s likelihood engines are able to provide accurate estimations of demographic parameters even for misspecified models. We improve model parameters for two empirical datasets of inbred species.

## Introduction

The evolutionary forces form a genetic variety of closely-related species and populations. Principal historical events like divergence, population size change, migration and selection could be reconstructed from the genetic data using different algorithmic and statistical approaches. Inference of complex demographic histories is widely applied in conservation biology studies to identify major events in the population’s past [1, 2, 3]. It supplements archaeological information about the historical processes that have left no paleontological records. Finally, demographic histories form the basis for subsequent population studies and medical genetic research.

In recent years many methods for demographic inference have appeared to investigate the demographic histories of species or populations [4, 5, 6, 7, 8, 9, 10, 11, 12]. Some of them give a point estimate for the unknown demographic parameters [4, 7, 11] while others give the distribution thereof [5, 6, 8]. In this paper we focus inclusively on the former. Most methods that provide point estimates consist of two independent components. The first component provides means to compute data statistics under a proposed demographic history and compare them with real data by the log-likelihood value. One of the most widely-used data statistics is the allele frequency spectrum (e.g., [4, 7, 10]). However, newer methods based on two-locus [13] and linkage disequilibrium (LD) statistics [14, 15] have also become available. This paper will denote the first component as an *likelihood engine*. The second component of existing tools is *optimization*. It requires a user-defined model of demographic history and performs a search of the maximal likelihood model parameters using different optimization algorithms. While a number of optimization techniques are provided, they often turn out to be ineffective in practical applications [16].

In 2020 we presented a new software GADMA [16] for unsupervised demographic inference from the allele frequency spectrum (AFS) data. It provides a common interface for various already existing likelihood engines and introduces new global search optimization based on a genetic algorithm. GADMA does not require complicated model specification. Instead, it takes *model structure* that determines how many time epochs are included in the model. Previously, models of demographic history were parameterized only by continuous parameters and had fixed population size dynamics. Constant size or exponential growth could be examples of such dynamics. GADMA’s model with structure extends the regular concept of a model by including dynamics as discrete model parameters. Thus, it can automatically construct history as a sequence of time epochs with desired parameter types from blocks of constant, linear and exponential size changes. The researcher has control over the types of model parameters to infer. For example, all migrations could be disabled.

It was shown that the proposed genetic algorithm approach in GADMA has better performance than previously existing optimization algorithms both on simulated and real datasets [16]. GADMA proved itself capable of finding demographic histories that attain higher log-likelihood than the histories reported in the literature and obtained using the default optimization routines of ∂a∂i or *moments*. Moreover, using demographic models with structure, GADMA was able to find a new demographic history of modern human populations that is both paleontologically plausible and has better log-likelihood than the existing “Out-of-Africa” scenario from Gutenkunst et al. [4]. Since its initial publication, GADMA has been applied in several studies on a variety of species: Xiong et al. [17], Valdez and D’Elía [18], Pazhenkova and Lukhtanov [19], Cassin-Sackett et al. [20], Buggiotti et al. [21].

The initial version of GADMA features only two likelihood engines: ∂a∂i [4] and *moments* [7]. Both of these engines compute the allele frequency spectrum statistics using Wright-Fisher diffusion-based approach and, thus, provide similar results. Among the variety of other available tools, we can highlight methods based either on AFS (*momi2, fastsimcoal2*), LD statistics (*momentsLD*), or haplotype data (diCal2) as potential additions to the supported engines in GADMA. Some already implemented features of ∂a∂i and *moments*, like the inference of selection and dominance rates, are not included in the first version of GADMA. Both ∂a∂i and *moments* have been upgraded since these programs were first published and since GADMA’s initial release. For example, ∂a∂i introduced inference of the inbreeding coefficients [22], started to support demographic histories involving four and five populations and enabled GPU support [23]. In light of these advancements, we have sought to extend GADMA in several directions to support new features and engines and further enhance its optimization algorithm.

In this paper, we describe new capabilities implemented in GADMA2. We compare supported likelihood engines of GADMA2 on two simulated datasets for different demographic models. Furthermore, we demonstrate the efficiency of the updated version on two empirical datasets of inbred species from Blischak et al. [22].

GADMA2 has an updated core codebase and implements a more efficient and flexible unsupervised demographic inference method. The improved version extends the initial GADMA in several ways (Figure 1). First, the genetic algorithm in GADMA2 is improved by hyperparameter optimization. New values of the genetic algorithm hyperparameters that provide more efficient and stable convergence are found. Second, GADMA2 provides more flexible control of the model specification for automatic model construction. For example, it is possible to include inferences about selection and inbreeding coefficients. Third, two new likelihood engines are integrated: *momi2* and *momentsLD*. Thus, GADMA2 supports four engines overall. Lastly, several functional enhancements are integrated, including the ability to use data in VCF format and new engines for history representation and visualization (*momi2* and *demes*).

**Figure 1.**
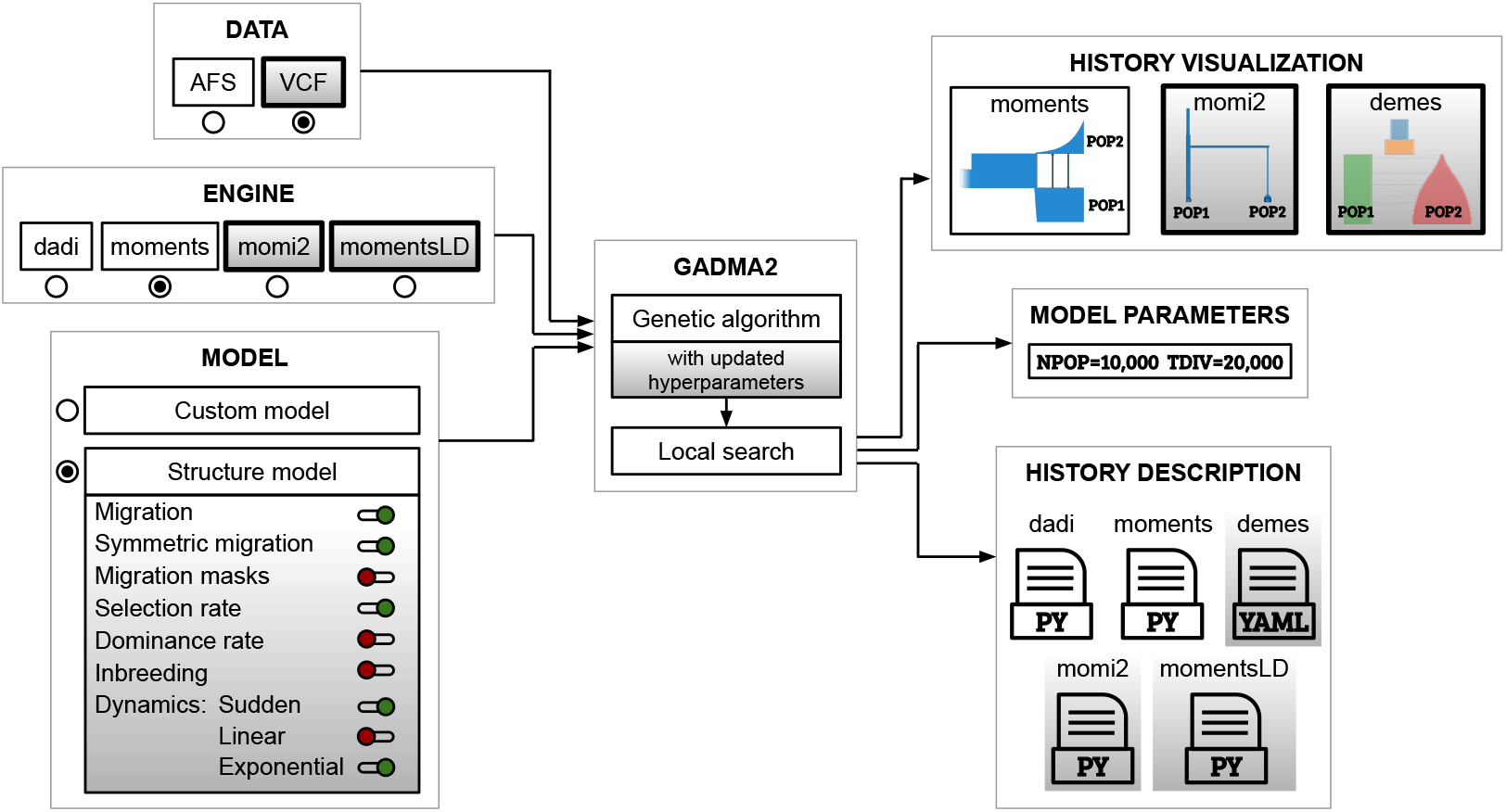
Scheme of GADMA2. New features and enhancements are marked with a gradient grey colour. GADMA2 takes input genetic data presented in either AFS or VCF formats, engine name and model specifications and provides inferred model parameters, visualization and descriptions of the final demographic history.

## Materials and methods

### Datasets

We use several datasets in this work. Datasets for the hyperparameter optimization are taken from the Python package `deminf_data v1.0.0` (Figure S1) that is available on GitHub via the link: https://github.com/noscode/demographic_inference_data. `Deminf_data` contains various datasets with both real and simulated AFS data. Simulations are performed with *moments* [7] software. Each dataset is named according to the convention described in Figure S1 and includes a) the allele frequency spectrum data; b) the model of the demographic history; and c) bounds of the model parameters. Full descriptions of the data and demographic model parameters of datasets are available in the repository on GitHub. For hyperparameter optimization we used ten datasets from `deminf_data`: four as training problem instances and six for testing. Basic descriptions of these datasets are available in section S1.1 of Supplementary Materials.

The performance of GADMA2’s engines is evaluated on two simulated datasets: populations of fruit flies and orangutan species. For simulation purposes we used a previously described scenarios available within the *stdpopsim* library [24]. Each dataset simulated by *msprime* engine [27] includes genetic data of five diploid individuals per each population.

Li and Stephan [25] presented the demographic history of *Drosophila melanogaster* populations from Africa and Europe. The visual representation of the history is shown in Figure 2a. The African population is characterized by a single instantaneous expansion. The origin of the European population is a result of the divergence of a very limited number of individuals followed by instantaneous expansion. Five autosomal chromosomes with a total length of 0.11 Gbp are simulated under this demographic history. We use mutation rate equal to 5.49 *·* 10^−9^ per base per generation [28] and recombination rate of 8.4 *·* 10^−9^ per base per generation [29].

**Figure 2.**
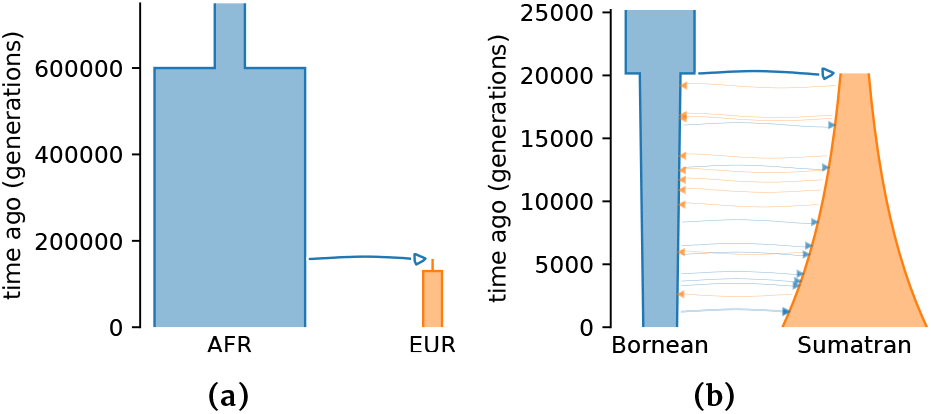
Demographic histories used in data simulations powered by *stdpopsim* [24] for performance comparison of GADMA2 likelihood engines. (a) History of African (AFR) and European (EUR) populations of *Drosophila melanogaster* from Li and Stephan [25]. (b) History of *Pongo pygmaeus* (Bornean) and *Pongo abelii* (Sumatran) orangutan species from Locke et al. [26].

The demographic history of the Bornean (*Pongo pygmaeus*) and Sumatran (*Pongoabelii*) orangutans was originallyinferred in Locke et al. [26] and is shown in Figure 2b. Specifically, it is an isolation-with-migration history that describes the ancestral population split followed by the exponential growth of Sumatran and an exponential decline of Bornean orangutans. We simulate 23 autosomal chromosomes with a total length of 2.87 Gbp. The mutation rate used in the simulation is equal to 1.5 *·* 10^−8^ per site per generation [30]. Averaged recombination rates for each chromosome are taken from the *Pongo abelii* recombination map inferred in Nater et al. [30].

Datasets for the demographic inference of inbred species are taken from the original paper Blischak et al. [22]. The 11 *×* 5 AFS data for two populations of the American puma (*Puma concolor*) was constructed on the basis of Ochoa et al. [31]. The AFS data for 45 individuals of domesticated cabbage (*Brassica oleracea*) was obtained from publicly available resequencing data [32, 33]. Both allele frequency spectra are folded due to a lack of information about ancestral alleles. Datasets are presented in the repository of the original article and are available via the following link: https://github.com/pblischak/inbreeding-sfs.

### Hyperparameter optimization

GADMA uses a genetic algorithm to optimize the demographic parameters [16]. A *hyperparameter* is usually defined as a parameter of an algorithm. The performance of any algorithm depends on its hyperparameters, and optimization of their values can significantly improve the overall efficiency. As an example of a hyperparameter, we can consider the number of demographic models in one iteration of the genetic algorithm. Several techniques can be used for the optimization of hyperparameters, and Bayesian optimization is one of the most popular methods [34]. The efficient method based on the Bayesian optimization is implemented in SMAC software [35, 36]. It has been applied in a number of studies including optimization of neural networks [37, 38, 39].

SMAC addresses the algorithm configuration problem, which involves determining the optimal hyperparameters for a given algorithm across multiple instances. As such, it needs to solve a multi-objective optimization problem. To do this SMAC uses a heuristic approach that optimizes a single objective function, the average value of the given cost function across the entire set of problem instances. Here we will refer to the objective function value as the *SMAC score*. The researcher usually selects the cost function according to the goal of the optimization process. This cost function is typically based on factors such as the time required to solve the problem or the quality of the solution achieved within a specific budget. It is important to exercise caution when working with SMAC score: since values of cost function are averaged across a given set of instances, it is essential that they have the same scale.

We use SMAC to tune the hyperparameters of the genetic algorithm in GADMA2. We divide all datasets into two groups of training and test datasets. The optimization is performed for GADMA’s genetic algorithm using *moments* engine and four training datasets as problem instances. Test datasets are used to validate the performance of the new configurations after optimization using SMAC. To ensure that the cost function has the same scale across all problem instances, we select the training datasets such that they have an identical number of populations and sample size. We choose the cost function as the best log-likelihood value achieved by GADMA’s genetic algorithm within a fixed number of likelihood evaluations. Regular GADMA’s pipeline, though, may require less or more evaluations to run depending on the dataset as it has a stop criterion that is based on the convergence. We take 200 times the number of dataset parameters as an allowed number of likelihood evaluations for the genetic algorithm runs in SMAC. Such a number of evaluations is a trade-off between speed and accuracy: according to the convergence plots, the convergence of default genetic algorithm optimization is slowing down at this point and is very close to the plateau walk (Figure S2, Figure S3). We note that we count the log-likelihood evaluations rather than the iterations of the genetic algorithm, as one iteration may involve multiple evaluations and the number of evaluations can vary across different configurations.

We perform several attempts of SMAC optimization for different variants of hyperparameter configurations. The descriptions and domains of all hyperparameters are given in Table S1 and Table 1 correspondingly. Each attempt took two weeks in 10 parallel processes (Intel® Xeon® Gold 6248). First, optimization of all genetic algorithm hyperparameters is executed. Then two discrete hyper-parameters (`gen_size` and `n_init_const`) are fixed to five manually picked combinations of domain values. SMAC is used to find optimal continuous hyperparameters for each combination. Four combinations were excluded from the analysis. Hyperparameter `gen_size` that corresponds to the size of generation in a genetic algorithm is not tested for the value of 100 due to relatively slow convergence. This eliminates three combinations. Additionally, the constant of initial design `n_init_const` equal to 5 is excluded for a case of `gen_size` equal to 50 as it provides a small number of solutions for the first generation.

**Table 1.** Hyperparameters of the genetic algorithm in GADMA, their domains used in SMAC and final values after each optimization attempt with SMAC. Hyperparameter values from attempt 1 are equal to the default GADMA values as SMAC failed to find a better configuration. For each attempt of 2-6 attempts two discrete hyperparameters (`gen_size` and `n_init_const`) are fixed in order to gain SMAC efficiency.

| Hyperparameter ID | Domain | Attempt number |  |  |  |  |  |
| --- | --- | --- | --- | --- | --- | --- | --- |
|  |  | 1 (default) | 2 | 3 | 4 | 5 | 6 |
| *gen_size* | {10, 50, 100} | 10 | 10* | 10* | 10* | 50* | 50* |
| *n_init_const* | {5, 10, 20} | 10 | 10* | 5* | 20* | 10* | 20* |
| *p_elitism* | [0, 1] | 0.20 | 0.30 | 0.30 | 0.30 | 0.40 | 0.40 |
| *p_mutation* | [0, 1] | 0.30 | 0.20 | 0.20 | 0.20 | 0.08 | 0.10 |
| *p_crossover* | [0, 1] | 0.30 | 0.30 | 0.30 | 0.30 | 0.42 | 0.46 |
| *p_random* | [0, 1] | 0.20 | 0.20 | 0.20 | 0.20 | 0.10 | 0.04 |
| *mutation_strength* | [0, 1] | 0.200 | 0.776 | 0.370 | 0.534 | 0.833 | 0.528 |
| *const_mutation_strength* | [1, 2] | 1.010 | 1.302 | 1.290 | 1.648 | 1.199 | 1.492 |
| *mutation_rate* | [0, 1] | 0.200 | 0.273 | 0.886 | 0.882 | 0.595 | 0.345 |
| *const_mutation_rate* | [1, 2] | 1.020 | 1.475 | 1.942 | 1.417 | 1.645 | 1.472 |

\*These values are fixed during the hyperparameter optimization with SMAC.

Overall, we make six attempts of hyperparameter optimization using SMAC. Unlike the training datasets, the additional six test datasets are selected to be diverse and, as a result, have varying scales of their likelihood functions. Therefore, it is not correct to compare new configurations obtained from different SMAC attempts using the SMAC score calculated across both training and test datasets. Furthermore, the genetic algorithm within the SMAC framework was terminated earlier than during a regular GADMA run. Nevertheless, we first validate the efficiency of SMAC and make preliminary comparisons of the new configurations for the AFS-based engines (*moments*, ∂a∂i and *momi2*) using SMAC scores evaluated across 128 independent runs. Then we compared new configurations by the log-likelihood values obtained from full genetic algorithm runs using the usual GADMA stop criterion. We use the same or equivalent criteria as the initial GADMA version to terminate the genetic algorithm [16]. For example, the genetic algorithm with a configuration with a generation size (`gen_size`) of 10 stops after 100 iterations without improvement, while the equivalent number of 20 iterations without improvement is used as a stop criteria for configurations with `gen_size` equal to 50.

The new configurations are compared as follows. First, we measure the speedup, which is the average fraction of the log-likelihood evaluations saved by a new configuration compared to the default configuration. Then, we compare each new configuration against the default configuration using the resulting likelihoods and determine its performance as *better, worse*, or *incomparable* for each dataset. For a fixed dataset, we consider a new configuration to be better if the median and both quartiles of the 128 likelihood values are higher than these quantities for the default configuration. If the median and both quartiles are lower, we consider the performance on the dataset to be worse. Otherwise, we declare the case incomparable. We aim to select a configuration taking into account both the speedup and the likelihood comparisons against the default configuration. Our goal is to find a configuration that is faster and performs better than the default configuration on as many datasets as possible, while also minimizing the number of datasets where it performs worse.

More information and details are available in section S1 of Supplementary Materials.

### Performance test of GADMA2 engines

Four engines supported by GADMA2 (∂a∂i, *moments, momi2, momentsLD*) are compared on two simulated datasets of fruit fly populations and orangutan species. For each dataset we test several models of the demographic history. The first two models are based on the ground truth history used in the simulations but differ in the presence of migration. Then we infer parameters for two structure models with and without migration using the GADMA2 feature for automatic demographic inference. For the orangutan dataset three additional models with pulse migrations are analyzed. The performance of all four engines is compared, however, *momi2* engine is not tested for models with continuous migration as it does not support it. We run GADMA2 inference eight times for each engine and model. Parameters of the history with the best log-likelihood are reported. Mutation and recombination rates for demographic inference are taken the same as in the data simulation. Their values are available in Datasets section.

Using GADMA2 engines we find and compare parameters for four models of *Drosophila melanogaster* demographic history (Table S5). Model DROS-NOMIG is an isolation model with instantaneous size change of the African population followed by separation of the European population which experiences two epochs of constant sizes. Population sizes during these epochs are not dependent with each other. Model DROS-MIG describes the identical to DROS-NOMIG scenario but includes continuous asymmetric migration between populations from their divergence till presence. Both models DROS-NOMIG and DROS-MIG align with the original isolation history used for data simulation. Lastly, we test two models with (DROS-STRUCT-MIG) and without (DROS-STRUCT-NOMIG) migration for structure (2, 1). This notation means a model consisting of two epochs before the ancestral population split followed by divergence and one epoch for each of the two subpopulations. More details on model structure specification can be found in Noskova et al. [16]. By their definition, these structure models are misspecified due to simplification of the European population’s history: a two-epoch scenario of the European population is approximated by one epoch with either constant size, linear or exponential change.

We analyze engines’ performance on the orangutan dataset for seven demographic models (Table S13). Model ORAN-NOMIG is isolation with the ancestral population split followed by the exponential size changes of the Sumatran and Bornean orangutans. Model ORAN-MIG aligns with the history used in data simulation and describes an isolation-with-migration with the ancestral population split followed by the exponential size changes of the Sumatran and Bornean orangutans. Additional two models with structure (1, 1) without (ORAN-STRUCT-NOMIG) and with continuous migration (ORAN-STRUCT-MIG) are included in the analysis. We note that the original history contains gene flow and can be correctly estimated using ORAN-MIG and ORAN-STRUCT-MIG models.

In order to overcome *momi2*’s limitation on continuous migrations presented in the orangutan history, we tested the engine for additional demographic scenarios with pulse migrations. A different number of pulse migrations with equal rates are uniformly distributed within the epoch between the present time and species divergence time. ORAN-NOMIG and ORAN-STRUCT-NOMIG models are compared with three additional demographic models: 1) with one pulse migration (ORAN-PULSE1), 2) with three pulse migrations (ORAN-PULSE3), and 3) with seven pulse migrations (ORAN-PULSE7).

### Inference of inbreeding coefficients

We perform demographic inference with GADMA2 using the data of the American pumas (*Puma concolor*) and domesticated cabbage (*Brassica oleracea* var. *capitata*) from Blischak et al. [22]. For each dataset parameters of two demographic models are inferred: 1) model from the original paper without inbreeding; 2) model from the original paper with inbreeding. Each demographic inference is run 100 times, and the history with the highest log-likelihood value is selected. Two result histories are compared with the likelihood ratio test [40] to investigate which history best fits the data.

First, we use the same parameter bounds to repeat the demographic inference from Blischak et al. [22] with GADMA2. We compare the results of 100 runs of GADMA2 with the same number of results received using ∂a∂i’s optimization techniques. Then we perform another round of demographic inference with GADMA2 using wider bounds of parameters.

Performance of GADMA2 is compared with performance of two optimization techniques from ∂a∂i within the same setup as in Blischak et al. [22]. We first reproduce 100 launches of a single ∂a∂i’s optimization as they were conducted in the original study and measure average number of evaluations and time of execution. Next, we run ∂a∂i’s optimization with restarts meaning that the optimization is restarted multiple times for each run and the best log-likelihood parameters are considered as the result. In order to balance computational costs, number of restarts is determined to match the average number of evaluations of GADMA2. Notably, if average ∂a∂i’s single optimization run requires *X* likelihood evaluations and average GADMA2 run involve *Y* evaluations, we compare GADMA2 run with run of ∂a∂i optimization with 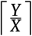 restarts. We consider the number of evaluations for comparison of computational costs. It is more reliable metric than the time of execution, as it is not affected by the specific hardware or parameter values used during optimization. Used optimization techniques from ∂a∂i require initial estimation of parameters which can be done by sampling from a wide range of distributions. To ensure correct comparison we use distribution from GADMA2 initial design to perform this initialization.

We report and compare the mean, the standard deviation and the best value of log-likelihood for 100 run repeats of GADMA2, of a single ∂a∂i’s optimization and of the ∂a∂i’s optimization with restarts. The optimization methods used for ∂a∂i runs are the BFGS algorithm [41, 42, 43, 44] for the American puma data and the BOBYQA method [45] for the domesticated cabbage data, as described in Blischak et al. [22].

Mutation rates, generation times and sequence lengths for parameter translation were taken from Blischak et al. [22]. Demographic parameters for *Puma concolor* are translated from the genetic to real units using a mutation rate of μ = 2.2 *×* 10^−9^, a generation time of 3 years, and a sequence length of 2,564,692,624 bp [31]. In the case of *Brassica oleracea* var. *capitata* population demographic parameters are translated using mutation rate of μ = 1.5 *×* 10^−8^, a generation time of 1 year, and a sequence length of 411,560,319 bp.

Godambe information matrix approach [40] was used for evaluation of the confidence intervals in the original paper [22]. This approach requires step size ϵ to estimate parameters’ uncertainty. The value of step size can influence the stability of Godambe approximation and several values should be tested to confirm consistent results between them. As in Blischak et al. [22] we estimate and compare uncertainties across a range of step sizes: : 10^−2^ – 10^−7^ by factors of 10. Reported confidence intervals for the final histories are estimated on 100 bootstrapped AFS data from the original paper using the Godambe information matrix with a step size equal to ϵ = 10^−2^ [40]. The scripts and data used for CI evaluation are taken from the repository of Blischak et al. [22] article: https://github.com/pblischak/inbreeding-sfs.

## Results and discussion

### Updated genetic algorithm

The genetic algorithm in GADMA2 is improved by the hyperparameter optimization implemented in SMAC software [35, 36]. Ten hyperparameters (Table 1) of the genetic algorithm were optimized during the first optimization attempt. SMAC performed 13,900 runs of the genetic algorithm and tested 2,222 different hyperparameter configurations. This process took two weeks of continuous computations on cluster. However, SMAC failed to find a better solution than the default one. We assume that such behavior may be caused by the presence of two discrete hyperparameters in the configuration. These hyperparameters are fixed to five specific combinations of the domain values during the next attempts of SMAC-based optimization of the remaining continuous hyperparameters.

As a result, we perform six attempts of hyperparameter optimization for different configurations of GADMA2, the result configurations are presented in Table 1. For each of these new configurations we manually evaluate the SMAC scores using 128 independent runs for each dataset and engine (*moments*, ∂a∂i and *momi2*). They can be found in Table S2 for the *moments* engine, Table S3 for the ∂a∂i engine and Table S4 for the *momi2* engine. The costs and results for ∂a∂i are very similar to those for *moments* supporting the idea that ∂a∂i and *moments* engines have very similar performance. Based on the obtained SMAC scores, the attempt 3 configuration is the best for *moments* and *momi2* engines and second best for ∂a∂i engine. However, we do not rely solely on the mean SMAC score as a selection criterion for these new configurations. This is because during SMAC runs the genetic algorithm was stopped earlier, and its full run performance may be different. Additionally, log-likelihoods between test datasets and between engines have different scales making direct comparison difficult.

To address this problem, we determine the best new configuration based on the performance of 128 full genetic algorithm’s runs using *moments* and *momi2* engines. For each configuration we measure the average speedup and indicate whether or not the result likelihood is better than for the default configuration. We do not perform the full runs for ∂a∂i engine due to high computational costs and its similarity to the *moments* engine. The boxplots of log-likelihood values and the required number of evaluations are presented in Figure S4 for *moments* engine and Figure S10 for *momi2* engine. The convergence plots of a genetic algorithm with different configurations on training and test datasets are presented in Figures S2, S3 for *moments* engine, Figures S6, S7 for ∂a∂i engine and Figures S8, S9 for *momi2* engine. The dataset counts for which new configurations demonstrate better, worse and incomparable performance comparing to the default configuration are presented in Figure S5 for *moments* engine and in Figure S11 for *momi2* engine.

On most datasets all new configurations require a smaller number of evaluations than the default genetic algorithm (Figures S4 and S10). There is only one dataset `2_ExpNoMig_5_Sim` (*moments* engine) for which the default configuration performs faster than all new configurations. In general, the configurations from attempts 3, 5, and 6 are the fastest. However, their log-likelihoods are worse than the default configuration on most datasets (Figures S5 and S11). Configurations from attempts 2 and 4 demonstrate best performance in terms of resulting likelihoods among new configurations. Moreover, genetic algorithm with hyperparameters from attempt 2 has better log-likelihood results for *moments* engine while the configuration from attempt 4 has better performance for *momi2* engine. Since the hyperparameter optimization used *moments* engine we choose the configuration from attempt 2 for the genetic algorithm in GADMA2. Figure 3 summarizes the improvement obtained by GADMA2 with the new hyperparameters as compared to the initial version. The new configuration saves around 10% of evaluations and provides better results on average compared to the default genetic algorithm. Some examples of the convergence plots that compare the previous version of genetic algorithm and the new version of genetic algorithm are presented in Figure 4.

**Figure 3.**
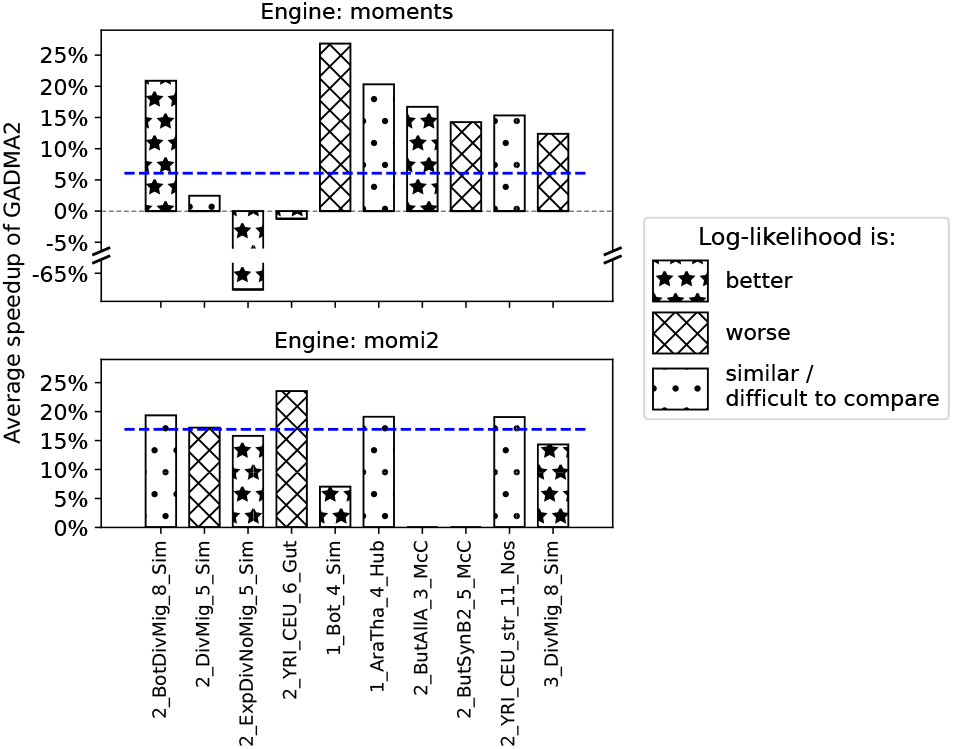
Performance comparison of the initial GADMA and GADMA2 with new hyperparameters. Bar size illustrate the average speedup of GADMA2 defined as the fraction of log-likelihood evaluations saved by the new version for each dataset. Blue dashed line demonstrates the average fraction of saved evaluations across all datasets. The bars’ hatching patterns indicate the improvement of the result log-likelihood based on median and quartiles. GADMA2 with new hyperparameters attains the average speed-up of 10% and provides better results on average comparing to the default configuration.

**Figure 4.**
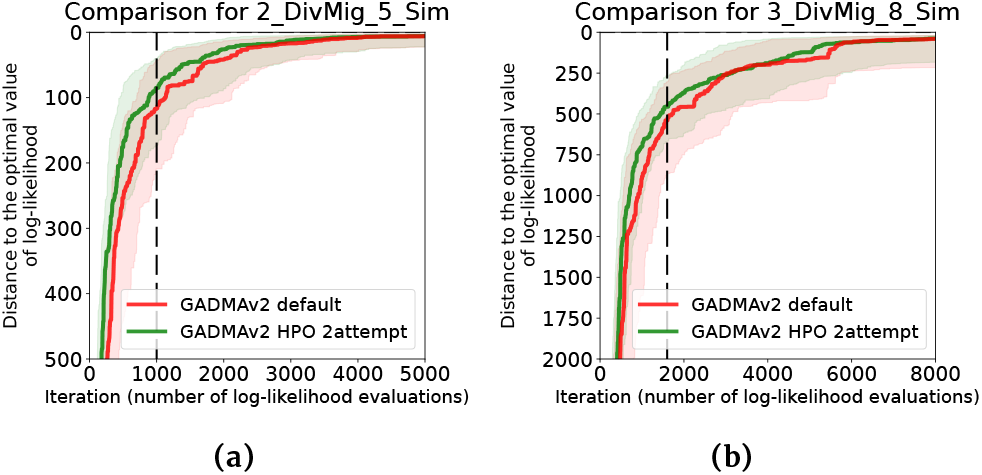
Example convergence plots for the default genetic algorithm configuration from the initial version of GADMA (red colour) and configuration obtained during attempt 2 of hyperparameter optimization with SMAC (green colour) on two datasets: (a) training dataset `2_DivMig_5_Sim`, (b) test dataset `3_DivMig_8_Sim`. For each configuration 128 independent optimization runs were performed. Solid lines correspond to median convergence over 128 runs and shadowed areas are ranges between the first (0.25) and third (0.75) quartiles. The vertical dashed black line refers to the number of evaluations used to stop a genetic algorithm in SMAC.

### Flexible structure model

Automatic demographic model construction is a central feature of GADMA. It replaces the fully manual choice of a model with a *model structure* specification. Traditionally, demographic models only have continuous parameters. Demographic structures, on the other hand, define the number of epochs before, after and between population splitting events and assign a discrete variable representing population dynamics type to each epoch. GADMA optimizes over these discrete variables alongside with the usual continuous ones, examining what would be a multitude of models in the traditional sense. GADMA2 gives the user more control over the search space in this setting.

#### Migration rates

One of the existing controls over model parameters is the opportunity to disable all migration events and to infer demographic history without any gene flow. GADMA2 now includes a new control handle to make migrations symmetric. Additionally, it allows for specific migrations to be disabled by setting up migration masks.

#### Selection and dominance rates

Both of the initially supported likelihood engines included in GADMA, ∂a∂i and *moments*, are able to infer selection and dominance rates. This inference approach first presented in Williamson et al. [46] assumes a single selection rate for the entire population while real genetic data could consist of regions with different rates. Despite this simplification such inference can provide useful estimations of selection. The first version of GADMA lacked the function to make these inferences and we have added these in the new version. GADMA2 enables the approximation of selection rates and dominance coefficients for automatically constructed demographic models.

#### Population size dynamics

GADMA2 provides additional flexibility for population size estimation during model construction. Previously, demographic parameters such as functions of population size changes were estimated within a fixed set of three possible dynamics: constant, linear, or exponential change. Now, the list of available population size dynamics in GADMA2 can be appointed to any subset of three basic functions. Thus, for example, linear size change can be excluded from the demographic inference if only constant and exponential dynamics are applicable, like in the case of *momi2* engine.

#### Inbreeding coefficients

Since the publication of the first version of GADMA, the supported likelihood engines were also upgraded. GADMA2 follows these changes and includes inference of inbreeding coefficients that were implemented in ∂a∂i [22]. Using this new feature included in ∂a∂i, we demonstrate that GADMA2 provides better and more stable results for inference of the demographic models obtained from data for the puma and cabbage reported by Blischak et al. [22] (Tables S21, S22 and Tables S26, S27).

### Data formats

Another improvement of ∂a∂i and *moments* is the ability to build an AFS dataset directly from a VCF file. Before this feature was implemented, this had to be done either manually or using another software like *easySFS* (https://github.com/isaacovercast/easySFS). GADMA2 is able to read data directly from a VCF file and downsize, exclude populations from, or build a folded AFS automatically. Such a feature allows broader and more convenient usage of GADMA2.

### New likelihood engines

In addition to ∂a∂i and *moments*, GADMA2 now includes two new likelihood engines: *momi2* [10] and *momentsLD* [14, 15]. Thus, four engines are provided in the common interface of GADMA2. Both ∂a∂i and *moments* engines are based on the Wright-Fisher diffusion and use allele frequency spectrum statistics for demographic inference.

*Momi2* implements a structured coalescent — backward-in-time stochastic process which is dual to the Wright-Fisher diffusion yet scales well to a large number of populations. It also uses AFS data as ∂a∂i and *moments*, but is computationally faster and can handle up to ten populations. However, *momi2* does not support continuous migration and linear change of population size.

Even though the allele frequency spectrum is one of the most popular statistics for demographic inference, it has limitations on how informative it can be [47]. The software *moments* has a submodule *momentsLD* dedicated to demographic inference using linkage disequilibrium (LD) statistics. In general, low-order two-locus LD statistics are used in *momentsLD*. A new likelihood engine using *momentsLD* is the first engine in GADMA that does not use AFS-based statistics.

Overall, GADMA2 now provides a choice of four likelihood engines and we encourage the community to extend this list.

### A new engine for demographic history representation

During demographic inference GADMA provides different textual and visual representations of the current best demographic history, such as generated Python code for all available likelihood engines or picture with visualized demographic history. Recently, a new Python package named *demes* [48] appeared to allow standard human-readable descriptions of demographic histories. GADMA2 includes *demes* as an engine to generate native descriptions and plots of demographic histories, which was only possible before using the *moments* or *momi2* engine. Figure 2 shows the examples of visual representations of demographic history using *demes*.

### Performance comparison of GADMA2 engines

We compare four likelihood engines supported by GADMA2 on two simulated datasets of fruit flies and orangutans. Several demographic models are used. Their description is provided in Performance test of GADMA2 engines section of Materials and methods.

The simulated parameter values of *Drosophila melanogaster* population history and their estimations inferred by engines in GADMA2 are presented in Tables 2, S7, S8 and S9 for DROS-NOMIG, DROS-MIG, DROS-STRUCT-NOMIG and DROS-STRUCT-MIG models correspondingly. The mean time of one log-likelihood evaluation and the mean number of evaluations averaged over inference runs are reported in Tables S9 and S10.

**Table 2.**
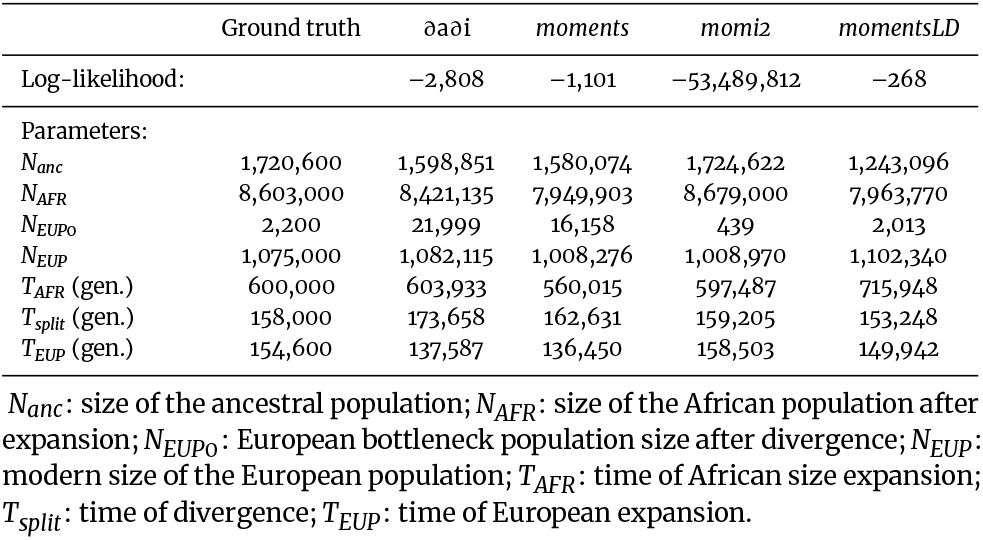
The demographic parameters of *Drosophila melanogaster* history without migration (DROS-NOMIG model) inferred with different engines in GADMA2. Ground truth are the parameter values from the paper Li and Stephan [25] used in simulation powered by *stdpopsim* [24]. Log-likelihood values are not comparable between different engines.

Estimations of orangutan history model parameters and their ground truth values are available in Tables S14, S15, S16 and 3 for ORAN-NOMIG, ORAN-MIG, ORAN-STRUCT-NOMIG and ORAN-STRUCT-MIG models respectively. The results of parameter estimations using *momi2* for models with zero, one, three and seven pulse migrations are presented in Table 4. The average time of one log-likelihood evaluation and the mean number of evaluations for used models and engines are reported in Tables S19 and S20.

**Table 3.** The demographic parameters of orangutan history with migration for structure (1, 1) (ORAN-STRUCT-MIG model) inferred with different engines in GADMA2. Ground truth is the simulated parameter values that were obtained from the original paper Locke et al. [26]. *Momi2* engine was excluded as it does not support continuous migrations. Log-likelihood values are not comparable between different engines.

| | Ground truth | $\partial\text{a}\partial\text{i}$ | *moments* | *momentsLD* |
| --- | --- | --- | --- | --- |
| Log-likelihood: |  | -1,220 | -1,106 | -53 |
| Parameters: |  |  |  |  |
| $N_{\text{anc}}$ | 17,934 | 17,925 | 17,854 | 17,685 |
| $N_{\text{Bor\_split}}$ | 10,617 | 10,432 | 10,498 | 10,529 |
| $N_{\text{Sum\_split}}$ | 7,317 | 7,492 | 7,355 | 7,155 |
| $N_{\text{Bor}}$ | 8,805<sup>exp</sup> | 9,282<sup>exp</sup> | 8,892<sup>exp</sup> | 8,592<sup>exp</sup> |
| $N_{\text{Sum}}$ | 37,661<sup>exp</sup> | 39,343<sup>exp</sup> | 37,443<sup>exp</sup> | 36,740<sup>exp</sup> |
| $m_{\text{Bor-Sum}} (\times 10^{-5})$ | 0.66 | 0.67 | 0.67 | 0.69 |
| $m_{\text{Sum-Bor}} (\times 10^{-5})$ | 1.10 | 1.07 | 1.09 | 1.13 |
| $T_{\text{split}} (\text{gen.})$ | 20,157 | 20,812 | 20,183 | 19,869 |

$N_{\text{anc}}$ : size of the ancestral population; $N_{\text{Bor\_split}}$ : size of *Pongo pygmaeus* at split; $N_{\text{Sum\_split}}$ : size of *Pongo abelii* at split; $N_{\text{Bor}}$ : modern size of *Pongo pygmaeus*; $N_{\text{Sum}}$ : modern size of *Pongo abelii*; $m_{\text{Bor-Sum}}$ : migration rate from *Pongo pygmaeus* to *Pongo abelii*; $m_{\text{Sum-Bor}}$ : migration rate from *Pongo abelii* to *Pongo pygmaeus*; $T_{\text{split}}$ : time of divergence. <sup>exp</sup> Exponential growth.

**Table 4.** The demographic parameters of orangutan histories without migration and with pulse migrations inferred using *momi2* engine in GADMA2. In ORAN-PULSE* models the time interval after divergence is divided into equal parts and pulse migrations are integrated between them. The inferred parameters show convergence to true values with an increase in pulse migration number. Ground truth is the simulated parameter values obtained from the original paper Locke et al. [26].

|  | Ground truth | NOMIG | STRUCT-NOMIG | Model ORAN-PULSE1 | PULSE3 | PULSE7 |
| --- | --- | --- | --- | --- | --- | --- |
| Number of pulse migrations | 0 (continuous) | 0 | 0 | 1 | 3 | 7 |
| Log-likelihood: |  | -48,541,453 | -48,545,934 | -48,437,315 | -48,391,684 | -48,377,617 |
| Parameters: |  |  |  |  |  |  |
| $N_{anc}$ | 17,934 | 19,331 | 19,086 | 19,220 | 18,461 | 17,997 |
| $N_{Bor\_split}$ | 10,617 | 6,187 | 8,453 | 8,731 | 8,715 | 10,086 |
| $N_{Sum\_split}$ | 7,317 | 7,719 | 11,668 | 4,165 | 5,412 | 6,409 |
| $N_{Bor}$ | 8,805 | 10,663 | 8,453 | 9,631 | 9,640 | 8,768 |
| $N_{Sum}$ | 37,661 | 54,184 | 49,595 | 59,929 | 43,123 | 38,030 |
| $m_{Bor-Sum}$ | $0.66 \times 10^{-5}$ | 0 | 0 | 0.065 | 0.057 | 0.025 |
| $m_{Sum-Bor}$ | $1.10 \times 10^{-5}$ | 0 | 0 | 0.206 | 0.084 | 0.036 |
| $T_{split}$ (gen.) | 20,157 | 11,270 | 11,668 | 16,211 | 20,086 | 20,809 |

$N_{anc}$ : size of ancestral population; $N_{Bor\_split}$ : size of *Pongo pygmaeus* at split; $N_{Sum\_split}$ : size of *Pongo abelii* at split; $N_{Bor}$ : size of *Pongo pygmaeus* after exponential decline; $N_{Sum}$ : size of *Pongo abelii* after exponential size change; $m_{Bor-Sum}$ : migration rate from *Pongo pygmaeus* to *Pongo abelii*; $m_{Sum-Bor}$ : migration rate from *Pongo abelii* to *Pongo pygmaeus*; $T_{split}$ : time of divergence in generations.

Below we present our general conclusions about the results. A more detailed comparison is available in section S2 of Supplementary Materials.

#### Fruit fly demographic history

Parameter values for models DROS-NOMIG and DROS-MIG that align with the ground truth are inferred accurately by all tested likelihood engines. Best estimations are obtained for the DROS-NOMIG model using *momi2* engine. The bottleneck European population size is approximated most accurately by *momentsLD* engine. Result histories for model DROS-MIG have worse values of log-likelihood than histories for the DROS-NOMIG model. Since DROS-NOMIG and DROS-MIG models are nested this indicates optimization failure. Nevertheless, they are able to catch general history and low migration rates. Thus, based on these results it is possible to assume population isolation and use further models without migrations for more accurate estimations.

We observe interesting results for the misspecified models with structure (2, 1). In the case of the DROS-STRUCT-NOMIG model, the ground truth history of *Drosophila melanogaster* is accurately approximated by *moments* and *momentsLD* engines only. The two-epoch history of the European population is approximated by exponential growth with a rate that differs between engines (Figure S12). The approximation made using *moments* engine aligns more closely with the actual history in terms of the mean population size and coalescent time, while the approximation from *momentsLD* is more accurate in terms of harmonic mean population size (section S2.1.1). We note that *momentsLD* engine also is able to provide similar history for the model DROS-STRUCT-MIG with migrations. However, ∂a∂i, *momi2* and *moments* for both models are hindered by the severe local optimum and were not able to achieve a global solution within eight GADMA2 runs. The alternative history is able to catch the European population history and low migration rates, yet, it does not reflect the instantaneous expansion of the ancestral population, and the parameter value for the African population size hits the upper bound. Using models with African population size fixed to the ancestral population size after expansion helps to overcome local optimum and achieve history similar to the ground truth (section S2.1.2 of Supplementary materials, Tables S11 and S12).

#### Orangutan demographic history

In the case of the orangutan simulated dataset all four engines provide similar demographic histories for the ORAN-NOMIG model without migrations. The predicted parameters are almost identical for ∂a∂i, *moments* and *momi2* which are AFS-based engines. Estimations for the modern sizes of populations are greater than the actual values used for the simulation. Moreover, the time of divergence is estimated to be lower: ∼12,000 vs. ground truth of ∼20,000. These discrepancies between predicted and simulated parameter values for the model ORAN-NOMIG could be explained by the fact that the model is oversimplified and lacks migration.

Model ORAN-MIG aligns correctly with the history used for data simulation. All tested engines provide estimations close to the simulated parameter values for the ORAN-MIG model.

The result demographic parameters for the ORAN-STRUCT-NOMIG model are close to the estimations obtained for the ORAN-NOMIG model. The population size dynamics are correctly inferred to be exponential for ∂a∂i and *momentsLD* engines. However, *momi2* and *moments* predict the constant size of the Bornean population. Although constant size approximates the Bornean population history relatively good, we demonstrate that our result is a consequence of the following model restriction. The model ORAN-STRUCT-NOMIG obliges the sum of Sumatran and Bornean population sizes after divergence to equal the ancestral population size. Ground truth history follows this rule, however, it is not fulfilled by the estimations inferred for the ORAN-NOMIG model. We ad-ditionally test the ORAN-NOMIG model with the same restriction on population sizes for *momi2* and *moments* engines. The best obtained scenarios have a worse log-likelihood value than histories with constant size of the Bornean population obtained for the ORAN-STRUCT-NOMIG model (section S2.2 of Supplementary materials).

Moreover, we remove the restriction on population sizes for ORAN-STRUCT-NOMIG model and infer parameters for the new modified model using *moments* and *momi2* engines. The history received for *momi2* engine is similar to those obtained for ORAN-NOMIG model and the history of Bornean population is estimated correctly by the exponential dynamic. However, even though *moments* engine also assumes exponential size change for Bornean population, it approximates the exponential growth of Sumatran population size by linear dynamic. Yet the history with linear approximation is similar to other histories obtained by *moments* for models ORAN-NOMIG and ORAN-STRUCT-NOMIG without migration. Furthermore, we ensure that such a model without the restriction but with linear size change is considered to be better than result history for ORAN-NOMIG model not only by *moments* engine but also by ∂a∂i and *momentsLD* (section S2.2 of Supplementary materials). Thus, we have observed that model misspecifications like absence of migrations may lead to confusions between exponential and linear dynamics but the results will still reflect the ground truth history.

The original demographic history of orangutan species used for data simulation is accurately reconstructed by ∂a∂i, *moments* and *momentsLD* engines within the ORAN-STRUCT-MIG model. Population size dynamics are inferred to be exponential for all tested engines. The parameters and values of log-likelihood are similar to the results for the ORAN-MIG model.

Finally, we analyze *momi2* engine performance for additional models ORAN-PULSE* with pulse events (Table 4). Pulse migration rates inferred by *momi2* differ significantly from continuous rates used in the simulation. However, it is important to note that pulse migration rates cannot be directly compared to continuous migration rates. As the number of pulse events increases, we expect the rates to decrease and it is supported by our results. For example, the migration rate from Bornean orangutans to Sumatran orangutans (*m*_*Bor*–*Sum*_) is inferred to be equal to 0.65 for model ORAN-PULSE1 with one pulse migration, to 0.057 for model ORAN-PULSE3 with three pulses and to 0.025 for model ORAN-PULSE7 with seven pulse events. It is crucial that other parameters converge to the simulated parameter values with an increased number of pulse events. Along these lines, population divergence time is estimated to be ∼11,000 generations for models ORAN-NOMIG and ORAN-STRUCT-NOMIG, ∼16,000 generations for the model ORAN-PULSE1 and ∼20,000 for models ORAN-PULSE3 and ORAN-PULSE7. The latter is close to the value of 20,157 used in the simulation. Parameter estimations for model ORAN-PULSE7 with seven pulse migrations are the most accurate among tested models. We assume the increase in pulse events number will lead to more accurate estimations yet require more computational resources. Thus, continuous migration is not supported in *momi2* engine but, to some degree, could be replaced by several pulse migration events.

### Usage case: inference of inbreeding coefficients

We use GADMA2 to reproduce demographic inference from Blischak et al. [22] for datasets of American pumas (*Puma concolor*) and domesticated cabbage (*Brassica oleracea* var. *capitata*). Blischak et al. [22] performed the demographic inference for two models without (model 1) and with inbreeding (model 2) using ∂a∂i’s optimization approaches.

First, we run GADMA2 with ∂a∂i engine and the same parameter bounds as in Blischak et al. [22] and compare the results of 100 repeats with the results obtained by ∂a∂i’s optimization techniques. The result statistics such as mean number of evaluations, mean execution time, mean and best values of log-likelihood are presented in Tables S21, S22 for American pumas and Tables S26, S27 for domesticated cabbage. They demonstrate that on average single GADMA2 run provides better and more stable results than single run of ∂a∂i’s optimization within 100 repeats. However, when optimization from ∂a∂i is restarted several times in order to match the computational costs of GADMA2 the results are not so consistent. In case of American puma populations final average and best log-likelihood values for GADMA2 are better than for ∂a∂i’s optimization with restarts. For domesticated cabbage inference optimization from ∂a∂i with restarts attains better average results than GADMA2. Yet number of restarts required to cover GADMA2 computational costs differs a lot between datasets and models and is always unknown in practice.

Several parameters of the result demographic histories obtained during first GADMA2 inference for both datasets received values close to their upper or lower bounds. In order to overcome this limitation, we perform another inference with wider bounds for parameter values and observe more reliable demographic parameters. The final values of the parameters and their confidence intervals are presented in Table S25 for American pumas and Table S30 for domesticated cabbage. Uncertainty estimates for confidence interval evaluation are consistent across different step sizes and are presented in Tables S23, S24 for American pumas and in Tables S28, S29 for domesticated cabbage. The visual representations of demographic histories using *demes* can be found in Figure 5 for American pumas and in Figure 6 for domesticated cabbage.

**Figure 5.**
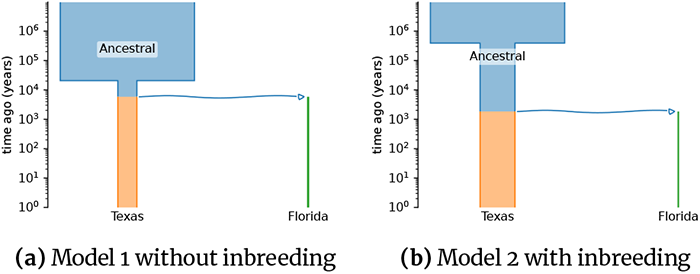
Demographic histories for Texas and Florida populations of American puma inferred with GADMA2. Figures are generated with the *demes* package [48]. Time is presented in a log scale.

**Figure 6.**
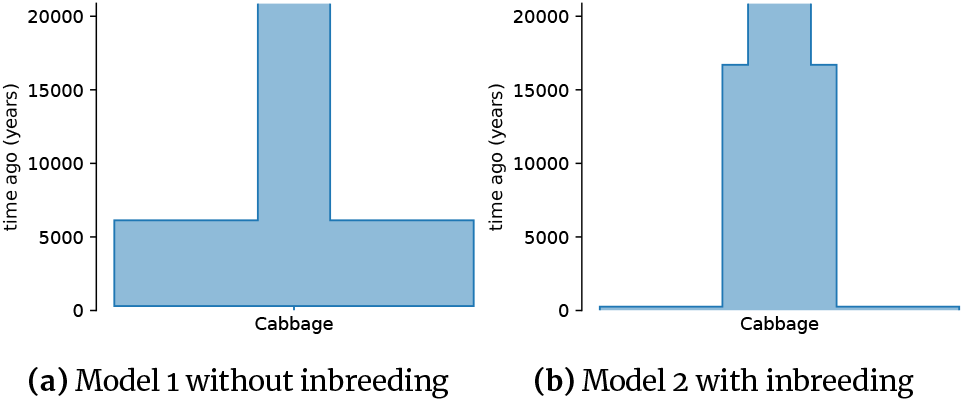
Demographic histories for a single population of domesticated cabbage inferred with GADMA2. Figures are generated with the *demes* package. In both models, the time of the most recent epoch is estimated to be small.

#### American puma demographic history

The best demographic histories obtained with GADMA2 have better values of log-likelihood (–452,492.70 vs –453,003.05 for model 1 and –316,115.56 vs. –318,058.08 for model 2) than those reported in Blischak et al. [22]. Similar values of population sizes are obtained except for the size of the Florida population which is estimated to be 860 and 374 individuals for model 1 and model 2 respectively compared to the 1,200 and 1,600 individuals estimated by Blischak et al. [22]. Time of divergence is estimated as 5,800 years ago for model 1 and as 1,800 for model 2. Inbreeding coefficients for model 2 are reported to be slightly higher than for the same model in Blischak et al. [22]: 0.453 for the Texas population and 0.628 for the Florida population. The Godambe-adjusted likelihood ratio statistic is 2,634.18 (P value = ∼0.0; Coffman et al. [40]), indicating that the model with inbreeding better describes data.

#### Domesticated cabbage demographic history

The best demographic histories obtained with GADMA2 for the domesticated cabbage population have better log-likelihood values (–24137.34 vs. –24330.40 for model 1 and –4267.32 vs. –4281.14 for model 2) than those than those received in Blischak et al. [22]. Values for the population sizes in the first and second epochs are inferred similar to the results from Blischak et al. [22]. However, the population size estimation for the most recent epoch in our results is lower (10 vs 592 individuals) for model 1 without inbreeding and higher (174,960,000 vs. 215,000 individuals) for model 2 with inbreeding than estimates obtained by ∂a∂i in [22]. The time duration of the epoch is also smaller for both models than estimated previously. In the case of model 1, the time parameter is very close to zero. The likelihood ratio test showed that the model with inbreeding better describes the data than the model without inbreeding (LRT statistic = 126.59, P value = ∼0.0; Coffman et al. [40]).

## Conclusions

GADMA2 is an extension of GADMA. It features an improved genetic algorithm, a more flexible automatic model construction setup and two additional demographic likelihood engines. We showcased GADMA2 by comparing different likelihood engines for two simulated dataset with various demographic models, including misspecified ones. Furthermore, we applied GADMA2 to infer demographic histories for two empirical datasets of inbred species, reporting updated parameters.

To improve the genetic algorithm we ran hyperparameter optimization powered by SMAC. We observed that discrete hyperparameters might hinder hyperparameter optimization, requiring much more iterations. Because of this, we manually picked five combinations of the discrete hyperparameters, running SMAC-based optimization of the remaining continuous ones for each fixed combination. We compared the set of optimal solutions on various datasets for three AFS-based likelihood engines of GADMA2: ∂a∂i, *moments* and *momi2*. We included the configuration that performed best when averaged across all likelihood engines as GADMA2’s new genetic algorithm. It is worth noting, however, that there was no one configuration that performed best for all of the likelihood engines simultaneously. We thus propose introducing engine-specific genetic algorithm configurations as a valuable direction for future work. Another prospective direction is population count-specific configurations, i.e. making the optimization of GADMA different for one, two and three populations. An important point here is that log-likelihoods for different datasets with a fixed population count are more comparable to each other. This could help SMAC’s heuristic target function better reflect the real multiobjective goal and thus improve its hyperparameter optimization performance.

GADMA’s automatic model construction setup was improved to allow forbidding specific migrations or making them symmetric, as well as restricting the admissible types of population size dynamics. Moreover, inference of selection and inbreeding coefficients was made possible in GADMA2. We note that the approach included in GADMA2 for inference of the selection and dominance rates is limited and can provide only simplified estimations. In order to perform more accurate inference of selection other approaches should be used [49].

Two new demographic likelihood engines, *momi2* and *momentsLD*, were incorporated into GADMA2. The former is based on a different mathematical model than ∂a∂i and *moments* and is computationally faster than them, but it does not support continu-ous migrations and linear population size growth. The latter is the first engine in GADMA2 that does not use allele frequency spectrum data for demographic inference and relies on linkage disequilibrium statistics instead. Furthermore, the new package *demes* was incorporated into GADMA2 as a representation engine providing textual and visual descriptions of demographic histories.

We analyzed the accuracy of GADMA2’s demographic likelihood engines on two simulated datasets: the dataset of fruit fly populations and the dataset of orangutan species. We used different demographic models, including models with structures. Some of these models align with the ground truth, while some are misspecified due to various simplifications. Similar performance was observed over all engines for the models that align with the ground truth. In this case, inferred demographic histories were close to the ground truth, and the types of population size dynamics were correctly recovered for models with structures.

Demographic inference with the misspecified models demonstrated interesting phenomena. For the misspecified models with structure and the fruit fly dataset, all the AFS-based engines were stuck at the same local optimum. However, the resulting demographic histories were still able to give some insights about the studied populations. The new LD-based engine *momentsLD* performed considerably better than the AFS-based engines. For the orangutan dataset both the AFS-based engines and *momentsLD* performed well. All slight discrepancies between estimated and ground truth values are consequences of models’ restrictions and misspecifications. It was found that sometimes exponential growth of population size could be misconstrued as linear growth with a similar rate for misspecified models that do not include migration events. Although the *momi2* engine does not support continuous migrations required to accurately model the ground truth, it performs well in approximating these with a number of pulse migrations. However, this approach is limited because larger numbers of pulse migrations increase computation time.

GADMA2 greatly simplifies performing such comparisons, the in-depth study of which seems a prospective work direction.

We reproduced the demographic inference setup of Blischak et al. [22] for the datasets of American pumas and domesticated cabbage. We compared performance of the fully ∂a∂i-based inference to GADMA2. GADMA2 attained higher log-likelihoods with lower variance across different runs than a single ∂a∂i optimization. GADMA2’s run times, however, are considerably longer than ∂a∂i’s, to the extent that ∂a∂i’s optimization can sometimes yield better results when restarted multiple times to match the computational costs of GADMA2. However, it may be difficult to determine the number of restarts needed, while GADMA2’s automatic termination makes for a simpler and, arguably, more reliable user experience.

Finally, we found updated parameters for models, both with and without inbreeding, for the datasets of American pumas and domesticated cabbage from Blischak et al. [22]. For each dataset, the best demographic histories include inbreeding. Our results, however, demonstrate very broad confidence intervals for some model parameters. The wide confidence intervals for the population size of domesticated cabbage during the most recent epoch can be explained by the fact that epoch length was inferred to be small, and very recent events are difficult to investigate with the

∂a∂i engine. However, the same results for the size of the Florida puma population and the population divergence time are difficult to explain. We only tested the demographic models from Blischak et al. [22], new models, however, can be built based on our results. GADMA2 extends the GADMA that has already shown itself as powerful and efficient software for the inference of complex demographic histories from genetic data. With its new application programming interface, GADMA2 can be easily improved further by integrating new likelihood engines, new optimization algorithms and automatic model construction routines.

## Availability of source code and requirements

GADMA2 is freely available from GitHub via the link https://github.com/ctlab/GADMA and can be easily installed via Pip or BioConda. Detailed documentation is located on the website https://gadma.readthedocs.io and includes user-manual, ready-to-use examples, and a section about Application Programming Interface (API). API enables an opportunity to use GADMA2 as a Python package and allows its optimization algorithms to be applied to any general optimization problem. An example of such usage is demonstrated for Rosenbrook function [50] optimization and is provided in the documentation.

- Project name: GADMA
- Version: 2.0.0
- Project home page: https://github.com/ctlab/GADMA
- Documentation: https://gadma.readthedocs.io
- RRID: RRID:SCR_017680
- biotoolsID: biotools:GADMA
- Operating system(s): Platform independent
- Programming language: Python
- Other requirements: Python3.6 or higher, other requirements are available within the documentation
- License: GNU GPL v3.

## Availability of Supporting Data and Materials

The scripts and the results of hyperparameter optimization experiments are saved in the repository and available via the link: https://github.com/noscode/HPO_results_GADMA. The results of GADMA runs for different hyperparameter configurations are stored as an archive available via the link: https://ctlab.itmo.ru/files/papers_files/GADMA2/comparison_on_datasets.zip. The results of experiments about inbreeding are added to the repository with the final demographic histories inferred in the original paper of GADMA and are located via the link: https://bitbucket.org/noscode/gadma_results.

