## Supplementary materials for "GADMA2: more efficient and flexible demographic inference from genetic data"

**References** **41**

### S1 Hyperparameter optimization

#### S1.1 Procedure

The performance of any algorithm depends on the values of its hyperparameters. The learning rate of the neural network is one of the classic examples of such dependence. One of the most popular frameworks for hyperparameters search is *Bayesian optimization*. The problem of finding hyperparameters that provide the best convergence not on one but several instances of data is called *algorithm configuration problem*. The algorithm configuration problem can be stated as follows: given a target algorithm $A$, a set of problem instances $I$ and a *cost function* (metric) $c$, seek for the best set of hyperparameters for $A$ on $I$ in regard to $c$. Usually, the cost function is based on the time required to solve the problem or on the solution's quality achieved within a given budget. SMAC (Hutter et al., 2011, Lindauer et al., 2022) is the software that implements Bayesian optimization for solving the algorithm configuration problem.

SMAC is based on Bayesian optimization, which is a model-based algorithm. It uses a *surrogate model* to estimate objective function and *acquisition function* to find new promising points for function evaluation. SMAC uses Random forest as a surrogate model for cost function approximation and expected improvement as an acquisition function to make predictions of promising hyperparameter configuration on each iteration. In contrast to classical Bayesian optimization, SMAC can handle several problem instances and therefore has some modifications. First, the objective function modeled with Random forest is a mean value of cost functions on instances. Another extension is the procedure of intensification. It is a mechanism that governs how to compare the new configuration with the existing best configuration (incumbent). Random Online Aggressive Racing (ROAR) implemented in SMAC considers a new configuration as a new incumbent if it is better than the current incumbent on the set of problem instances and random seed pairs. This set is extended each time a new configuration turns out to be worse than the incumbent. Such an algorithm considers that this set is always growing, and more and more comparisons are required to beat the current best configuration.

In summary, SMAC is an iterative algorithm that keeps the best-found configuration of hyperparameters. The new promising configuration is chosen and compared to the current incumbent within the intensification procedure in each iteration. The value of the cost function averaged over problem instances manages the comparison, as a result the best-by-average algorithm performance hyperparameter configuration is provided. We used SMAC to investigate hyperparameter values of the genetic algorithm in GADMA.

The optimization method, such as the genetic algorithm, has several hyperparameters, and their number varies within specific implementations. The genetic algorithm presented in GADMA maintains a set of problem solutions called generation. Each solution is presented as an array of values for objective function parameters, i.e., parameters of the demographic history. The initial generation is formed by the *initial design* procedure: a set of random solutions is created. The size of this set is determined by the value of hyperparameter `n_init_const`: the number of solutions in the initial generation is equal to the number of target parameters multiplied by `n_init_const`. The size of each generation in the genetic algorithm is equal to the value of the `gen_size` hyperparameter. The new generation is constructed iteratively with the help of mutation, crossover and selection of best by the value of likelihood models. The fractions of most adapted, mutated, crossed and random models that form a new generation are determined by `p_elitism`, `p_mutation`, `p_crossover`, `p_random` hyperparameters correspondingly. The special case is the mutation process:

Table S1: Short descriptions of GADMA genetic algorithm (GA) hyperparameters.

| Hyperparameter ID | Hyperparameter description |
|---|---|
| gen_size | Number of solutions in generation of genetic algorithm |
| n_init_const | Constant that determines number of random solutions that are created during initial design at the beginning of GA |
| p_elitism | Fraction of the best solutions that are taken to new generation |
| p_mutation | Fraction of mutated solutions in a new generation |
| p_crossover | Fraction of crossed solutions in a new generation |
| p_random | Fraction of random solutions in a new generation |
| mutation_strength | Initial parameter change probability for mutation |
| const_mutation_strength | Constant to change a mutation_strength during genetic algorithm according to one-fifth rule |
| mutation_rate | Initial rate of a parameter change during mutation |
| const_mutation_rate | Constant to change the mutation_rate during genetic algorithm according to one-fifth rule |

it is determined by mutation_strength and mutation_rate that define how many parameters of the model and how strong their values will change. Each of these two hyperparameters is changed during the genetic algorithm performance according to the one-fifth rule: the closer to the optimum we are, the smaller changes during the mutation process are. The constants of one-fifth rule (const_mutation_strength, const_mutation_rate) are additional two hyperparameters of the genetic algorithm. In total, we highlight ten hyperparameters for the genetic algorithm implemented in GADMA: two have integer values, and eight are continuous. The short descriptions of each hyperparameter are presented in Table S1.

SMAC uses a cost function averaged over problem instances as a target function to compare different configurations. The cost function can be based on either the runtime required to solve the problem or the solution quality achieved within a given budget. We follow the latter criteria to build a cost function for GADMA's hyperparameter optimization with SMAC. We use the log-likelihood achieved within a fixed budget as a metric that determines the quality of the configuration. The budget is measured in units of evaluations to be independent of specific hardware or fluctuations in the amount of time needed to perform one evaluation. The number of evaluations required by the genetic algorithm to converge is not always proportional to the number of parameters in a dataset. However, to ensure that the genetic algorithm has sufficient time to optimize larger datasets, we set the budget available for the algorithm as a linear function of the number of parameters in the dataset. Specifically, we choose the budget to be 200× the number of parameters as a compromise between the speed and accuracy of the genetic algorithm. Thus, for each configuration SMAC runs the genetic algorithm on training datasets for a fixed number of evaluations and takes the average best log-likelihood value as the result cost function of this configuration. We note that it is important to have comparable and well-balanced values of log-likelihood across training datasets to get accurate results when using a cost function averaged over problem instances.

The initial values of hyperparameters in the first version of GADMA were obtained manually within the demographic inference of two populations of modern humans for the model and data from Gutenkunst et al. (2009) (2_YRI_CEU_6_Gut dataset). Their values are presented in Table 1.

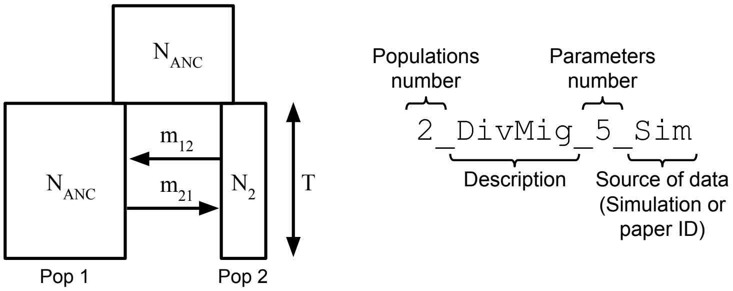

Figure S1: The model of the demographic history and naming convention of the example dataset from the `deminf_data v1.0.0` package.

Four datasets are chosen as training problem instances for SMAC. The result configurations are tested and compared to the default genetic algorithm on training and additional six test datasets. All used datasets are taken from Python package `deminf_data v1.0.0`. Each dataset presented in Table S2, Table S3 and Table S4 has a structured name (Figure S1) that is the sequence of: a) population number, b) a short description of the demographic model, c) a number of parameters, and d) information about the source of AFS data.

Each of four training datasets has demographic model and data for two populations: three simulated AFS data (`2_BotDivMig_8_Sim`, `2_DivMig_5_Sim`, `2_ExpDivNoMig_5_Sim`) and one real data (`2_YRI_CEU_6_Gut`) for modern human populations from Gutenkunst et al. (2009). Thus, datasets require similar resources for likelihood evaluations and are well-balanced for optimization with SMAC.

Test datasets are chosen to be more diverse: two datasets (`1_Bot_4_Sim`, `1_AraTha_4_Hub`) for one population with simulated and real data from Huber et al. (2018), two datasets (`2_ButAll_3_McC`, `2_ButSynB2_5_McC`) for two populations of butterflies from McCoy et al. (2014); one simulated dataset for three populations (`3_DivMig_8_Sim`); and one dataset (`2_YRI_CEU_struct_11_Nos`) with AFS data for two populations of modern human from Gutenkunst et al. (2009) and extended demographic history with structure (2, 1) that was observed in the original paper of GADMA Noskova et al. (2020). The notation of the model with structure (2, 1) means a model consisting of two epochs before the ancestral population split followed by divergence and one epoch for each of the two subpopulations. The dynamics of populations size changes are available for the inference within this model.

Six attempts of SMAC optimization are performed. During the first run of SMAC, all ten hyperparameters (Table 1) are optimized. During next five attempts two discrete hyperparameters (`gen_size` and `n_init_const`) are fixed to several values. During the second attempt of SMAC optimization discrete hyperparameters `gen_size` and `n_init_const` are fixed to default values (10 and 10 correspondingly) (Table 1). For the third and fourth attempts we test two alternative values (5 and 20) of the initial design constant (`n_init_const`). Then the number of demographic models on each generation of genetic algorithm (`gen_size`) is increased up to 50 and constant for initial design (`n_init_const`) is tested for values of 10 and 20. The value of `n_init_const` equal to 5 was excluded from the experiments as it provides a small number of solutions for the first generation that should be of size 50.

#### S1.2 Comparison of the new configurations

First attempt of SMAC optimization failed to find better configuration than the default one. We have obtained five new configurations for GADMA genetic algorithm from attempts 2-6 and compare them to the default configuration using several approaches.

First, average scores used in SMAC are independently computed for all configurations. SMAC rates configurations based on the cost function which is the mean log-likelihood attained within a fixed budget of evaluations averaged over all training datasets. These SMAC scores demonstrate the success of hyperparameter optimization yet provide a preliminary comparison of final configurations as the overall convergence of the genetic algorithm may be different from the convergence within a fixed budget. We compute SMAC scores for each dataset independently from 128 runs of early terminated genetic algorithm using *moments*, ∂a∂i and *momi2* engines (Tables S2, S3 and S4 respectively) We also report scores averaged across either training or test datasets and total scores averaged across all datasets. However, we note that these scores should be treated carefully as log-likelihoods have different scales between test datasets.

The convergence plots are presented in Figures S2, S3 for *moments* engine, in Figures S6, S7 for ∂a∂i engine and in Figures S8, S9 for *momi2* engine.

Incumbents for attempts 2, 3 and 4 where `gen_size` was fixed to the value of 10 show better SMAC scores averaged across either training, test or all datasets than the default configuration for all engines. In contrast, incumbents for attempts 5 and 6 with `gen_size` fixed to the value of 50 provide scores that are worse than those obtained for the default configuration in some cases. The attempt 3 configuration shows the best total mean score for *moments* and *momi2* engines. In case of ∂a∂i it is next to the the best and the configuration from attempt 4 provides the highest total score.

The final configuration is selected based on the comparison of accuracy and speed, considering 128 full runs of the genetic algorithm for each configuration and dataset using both the *moments* and *momi2* engines. Engine ∂a∂i is not analyzed due to its high computational costs and its similarity to *moments* engine. We report and compare the speed and log-likelihood accuracy results for each configuration. Figure S4 shows boxplots of the resulting log-likelihoods and the required number of evaluations for datasets received using *moments* engine. Figure S10 demonstrates the same plots for *momi2* engine.

To begin, we measure the speedup — the fraction of log-likelihood evaluations saved by new configuration compared to the default configuration. In order to compare log-likelihood accuracy of the configurations from attempts 2-6 to the default configuration, we categorize their performance into three groups: 1) better, 2) worse, and 3) undefined. The better category includes datasets where the new configuration shows higher median and quartiles of the resulting log-likelihoods compared to the default configuration. We consider the performance to be worse when both the median and quartiles are lower. Otherwise, we declare the performance to be incomparable. The histograms that demonstrate counts of datasets falling into each category for each configuration are presented on Figures S5 and S11 for *moments* and *momi2* engines correspondingly. We select configuration that is faster and performs better or similar (incomparable) than the default configuration on as many datasets as possible.

The speedups of the configurations from attempts 2-6 are for 6%, 30%, 11%, 28%, 51% *moments* engine and 16%, 35%, 15%, 37%, 55% for *momi2* engine. The total average speedups of configurations from attempts 2-6 compared to the default configuration are 10%, 32%, 13%, 32%, 53% respectively. Most new configurations require lesser number of evaluations than the default genetic algorithm, however, some of them loose accuracy due to fast convergence. We highlight configurations from attempts 3, 5 and 6 as such examples.

Among all new configurations, incumbents from attempts 2 and 4 show better log-likelihoods on a greater number of datasets and worse log-likelihoods on fewer datasets compared to the default configuration (Figures S5 and S11). When using the *moments* engine, the incumbent from attempt 2 performs the best, showing better log-likelihoods on 4 datasets and worse log-likelihoods on 2 datasets. The configuration from attempt 4 takes the second place, with 3 datasets demonstrating better log-likelihoods and 3 datasets with worse log-likelihoods. On the other hand, when using the *momi2* engine, the attempt 4 configuration demonstrates the best results. It showcases better log-likelihoods on 5 datasets and worse log-likelihoods on 2 datasets, surpassing the attempt 2 configuration, which has 3 datasets with better log-likelihoods and 2 datasets with worse log-likelihoods

We narrow our set of candidate configurations down to the configurations from attempts 2 and 4. On average attempt 4 configuration (13% speedup) was faster than incumbent from attempt 2 (10% speedup). Configuration from attempt 2 shows better log-likelihood results than configuration from attempt 4 for *moments* engine. Since the hyperparameter optimization with SMAC used *moments* engine, the configuration from attempt 2 is taken as a new hyperparameters of the genetic algorithm in GADMA2.

Based on our results we chose incumbent from attempt 2 as new hyperparameters of the genetic algorithm in GADMA2 but we note that attempt 4 configuration is the better choice when *momi2* engine is used.

##### S1.3 SMAC costs and convergence plots using *moments* engine

Table S2: Mean log-likelihood values (128 runs) for final configurations of six SMAC attempts on training and test datasets. Genetic algorithm was stopped at the same number of evaluations used in SMAC. Log-likelihood was evaluated with *moments* engine. Mean cost value on training datasets presented in the table is SMAC score that was used by SMAC intensification procedure. For attempt 1 SMAC failed to find better configuration than the default one. Best mean values are marked bold.

| Dataset | attempt number | | | | | |
|---|---|---|---|---|---|---|
| | 1 (default) | 2 | 3 | 4 | 5 | 6 |
| Mean cost on train datasets | −2,074.44 | −1,860.98 | **−1,798.29** | −1,843.54 | −1,990.94 | −2,006.94 |
| training datasets: | | | | | | |
| `2_BotDivMig_8_Sim` | −2,707.51 | −2,411.40 | −1,967.04 | −2,197.83 | −2,085.60 | **−1,963.56** |
| `2_DivMig_5_Sim` | −1,497.13 | **−1,439.16** | −1,523.42 | −1,453.18 | −1,471.58 | −1,441.97 |
| `2_ExpDivNoMig_5_Sim` | −2,936.62 | **−2,353.94** | −2,566.71 | −3,137.09 | −3,258.73 | −3,137.09 |
| `2_YRI_CEU_6_Gut` | −1,156.50 | −1,139.41 | −1,144.00 | −1,138.40 | −1,147.85 | **−1,134.40** |
| Mean cost on test datasets | −2,381.82 | −2,279.46 | −2,299.86 | −2,290.99 | −2,284.17 | **−2,268.37** |
| Test datasets: | | | | | | |
| `1_Bot_4_Sim` | −213.92 | **−193.49** | −212.19 | −194.40 | −196.82 | −212.42 |
| `1_AraTha_4_Hub` | −96.12 | **−93.05** | −96.45 | −94.03 | −95.04 | −95.53 |
| `2_ButAllA_3_McC` | −298.32 | **−290.30** | −300.36 | −294.21 | −306.68 | −293.99 |
| `2_ButSynB2_5_McC` | −216.84 | −214.78 | −216.51 | −214.53 | −214.93 | **−213.85** |
| `2_YRI_CEU_str_11_Nos` | −1,165.02 | −1,164.41 | −1,163.55 | −1,157.04 | −1,157.15 | **−1,150.26** |
| `3_DivMig_8_Sim` | −12,300.70 | −11,742.43 | −11,797.04 | −11,791.72 | −11,734.39 | **−11,637.73** |
| Mean cost on all datasets | −2,258.86 | −2,112.07 | **−2,099.23** | −2,112.01 | −2,166.88 | −2,163.80 |

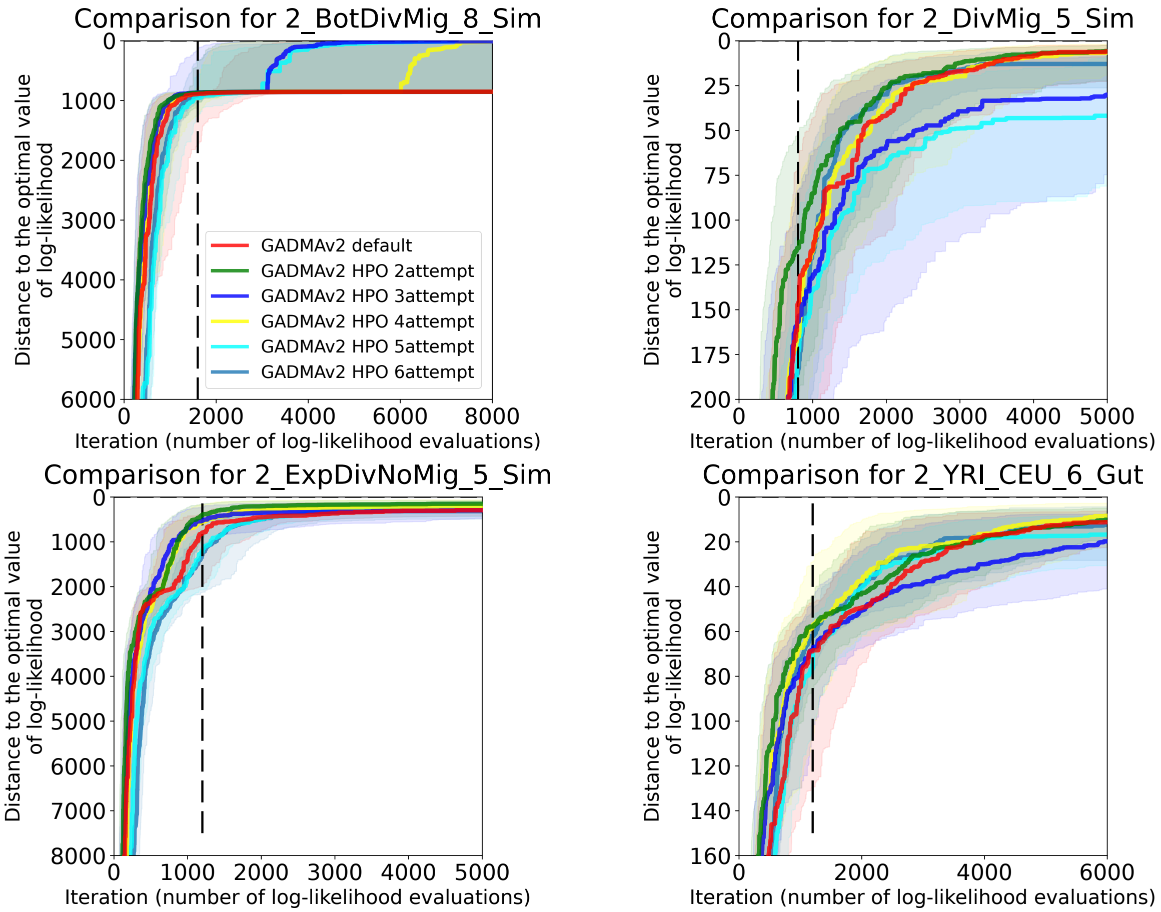

Figure S2: Convergence plots for six genetic algorithm configurations using *moments* engine on four training datasets: 1) the default genetic algorithm from the initial version of GADMA, 2)-6) configurations obtained during attempts 2-6 of hyperparameter optimization with SMAC. The abscissa presents the log-likelihood evaluation number, the ordinate refers to the distance to the optimal value of log-likelihood. Solid lines correspond to median convergence over 128 runs and shadowed areas are ranges between first (0.25) and third (0.75) quartiles. The vertical dashed black line refers to the number of evaluations used to stop a genetic algorithm in SMAC. The default configuration (red) and two configurations from attempt 2 (green) and attempt 6 (blue) were compared in terms of convergence on a greater number of iterations. The configuration from attempt 2 shows faster convergence on first iterations, the configuration from attempt 6 turns out to have better convergence at last iterations on three of four datasets.

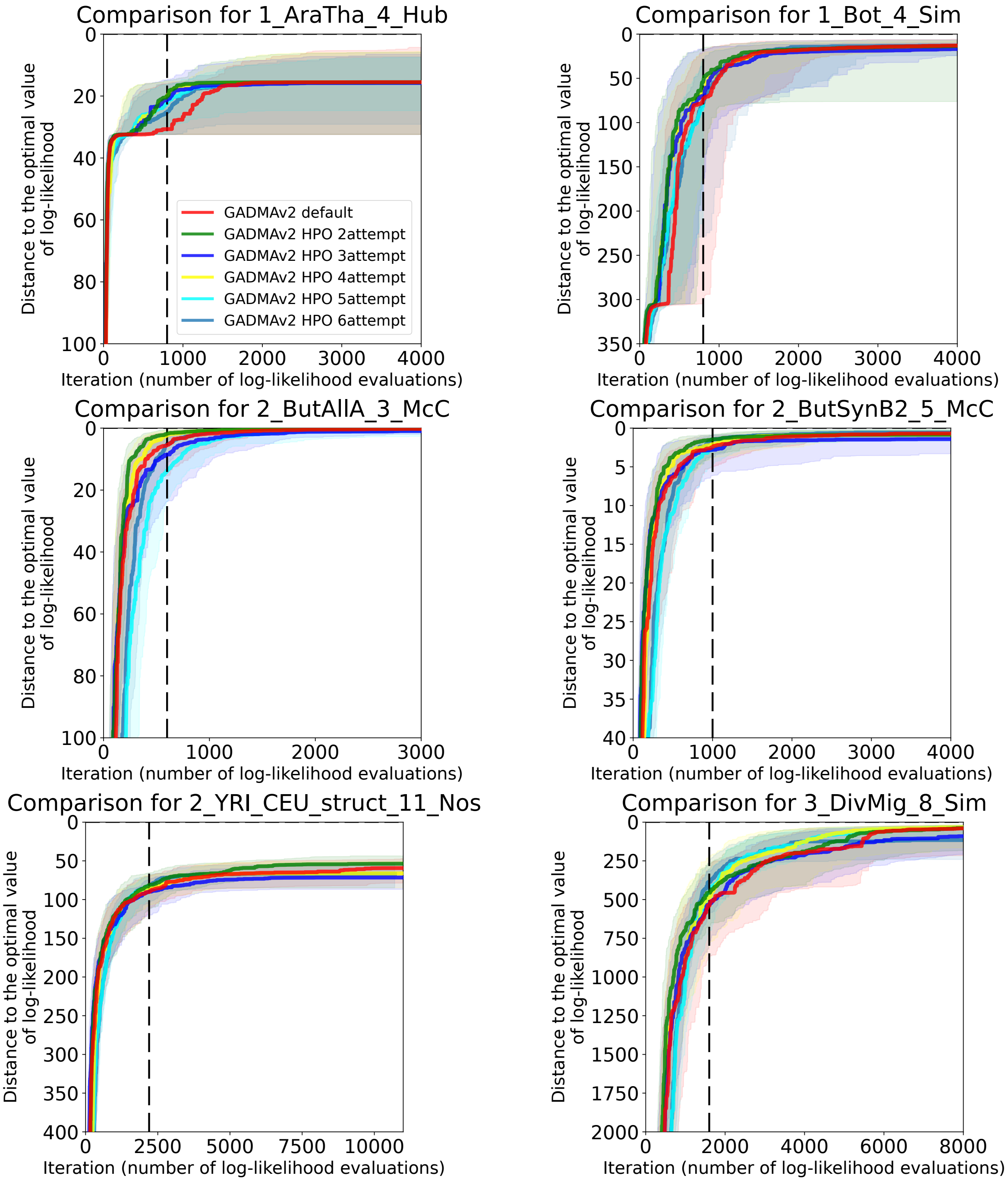

Figure S3: Convergence plots for six genetic algorithm configurations using *moments* engine on six test datasets: 1) the default genetic algorithm from the initial version of GADMA, 2)-6) configurations obtained during attempts 2-6 of hyperparameter optimization with SMAC. The default configuration (red) and two configurations from attempt 2 (green) and attempt 6 (blue) were compared in terms of convergence on a greater number of iterations. The configuration from attempt 2 shows faster convergence on first iterations, the configuration from attempt 6 turns out to have better convergence at last iterations on two of six datasets.

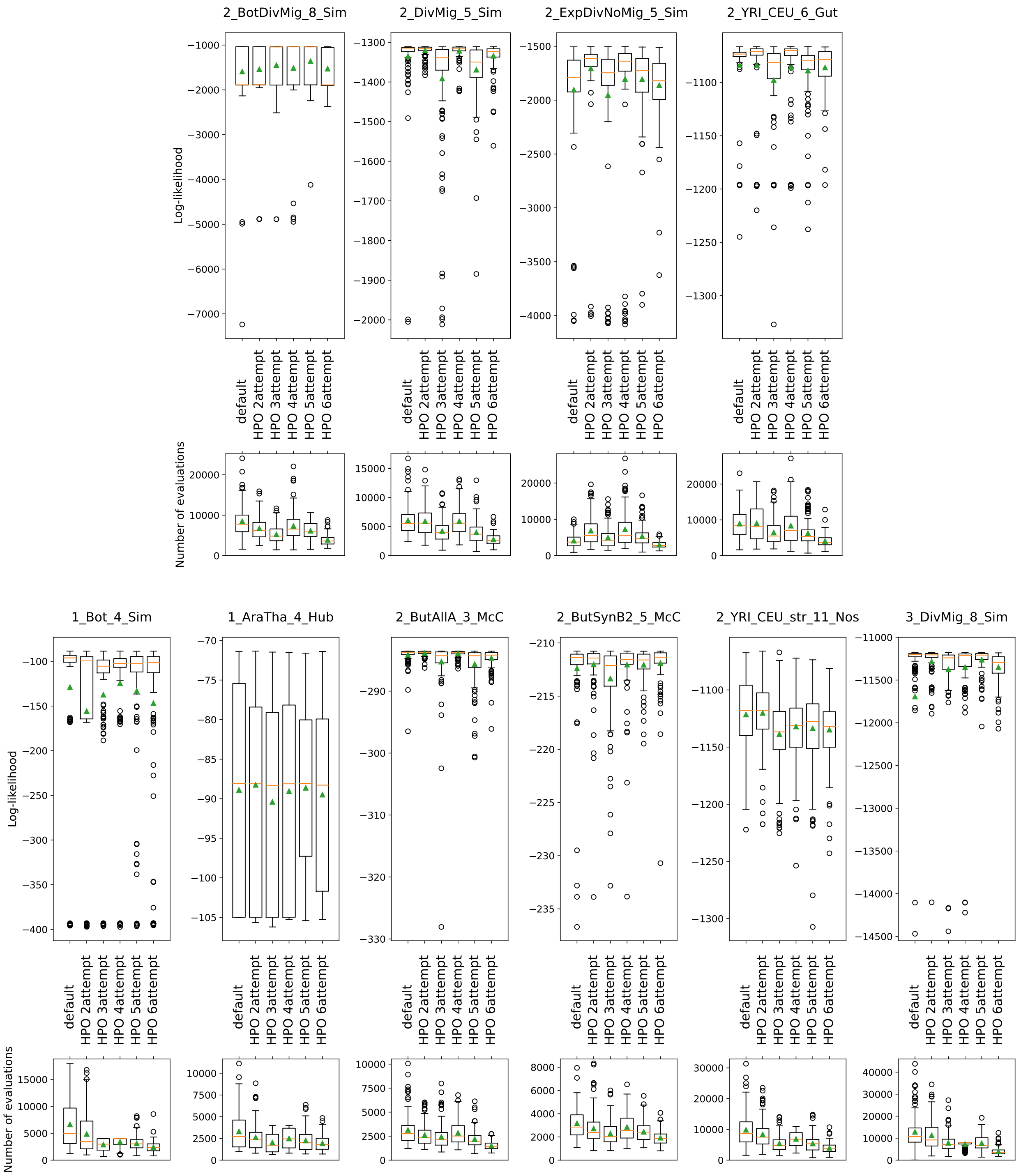

Figure S4: Boxplots of the eventual log-likelihoods and number of evaluations required for full runs of genetic algorithm with six configurations using *moments* engine. For each dataset two plots are presented: 1) the top plot shows distribution of 128 resulting log-likelihood values; 2) the bottom plot corresponds to the distribution of the evaluations' number required for genetic algorithms to terminate. Orange line on boxplot refers to the median value, greed triangle demonstrates the mean value.

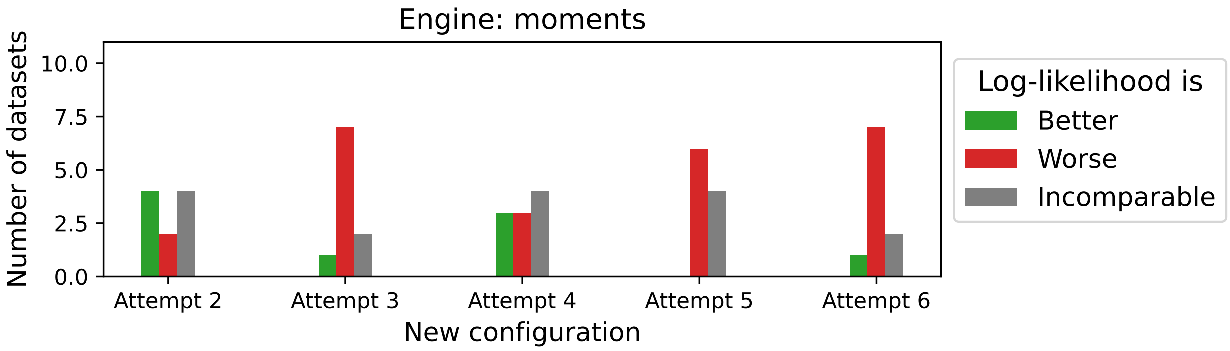

Figure S5: For each dataset, the performance of the new configurations from attempts 2-6 is categorized as either better, worse, or incomparable to the performance of the default configuration when using the *moments* engine. The histogram illustrates the count of datasets falling into each category for each configuration. Performance is considered to be better if both the median value and both quartiles of log-likelihoods are higher than those of the default configuration. Conversely, if both the median and quartiles are lower, the dataset is categorized as worse. Otherwise, the comparison is considered to be similar or undefined.

#### S1.4 SMAC costs and convergence plots using ∂a∂i engine

Table S3: Mean log-likelihood values (128 runs) for final configurations of six SMAC attempts on training and test datasets using ∂a∂i simulation engine. Genetic algorithm was stopped at the same number of evaluations used in SMAC. Best mean values are marked bold. Log-likelihood values and results are similar to *moments* engine.

| Dataset | attempt number | | | | | |
|---|---|---|---|---|---|---|
| | 1 (default) | 2 | 3 | 4 | 5 | 6 |
| Mean cost on training datasets | −1,939.03 | −1,848.47 | −1,805.90 | **−1,790.88** | −2,043.95 | −1,959.65 |
| training datasets: | | | | | | |
| 2_BotDivMig_8_Sim | −2,392.63 | −2,264.38 | −2,140.06 | −2,115.68 | −1,998.72 | **−1,908.14** |
| 2_DivMig_5_Sim | −1,494.41 | **−1,457.52** | −1,495.68 | −1,461.03 | −1,481.33 | −1,465.06 |
| 2_ExpDivNoMig_5_Sim | −2,720.98 | −2,534.38 | **−2,448.77** | −2,450.89 | −3,471.70 | −3,329.37 |
| 2_YRI_CEU_6_Gut | −1,148.08 | −1,137.60 | −1,139.10 | −1,135.92 | −1,284.13 | **−1,134.62** |
| Mean cost on test datasets | −2,321.25 | −2,288.71 | −2,295.75 | −2,280.61 | −2,283.11 | **−2,269.74** |
| Test datasets: | | | | | | |
| 1_Bot_4_Sim | −231.34 | −209.55 | **−170.73** | −192.93 | −216.42 | −228.53 |
| 1_AraTha_4_Hub | −93.27 | **−92.86** | −95.80 | −94.21 | −95.96 | −95.82 |
| 2_ButAllA_3_McC | −296.52 | **−291.90** | −301.06 | −297.96 | −301.25 | −298.50 |
| 2_ButSynB2_5_McC | −204.13 | −204.90 | **−191.29** | −214.57 | −216.21 | −192.83 |
| 2_YRI_CEU_str_11_Nos | −1,173.82 | −1,166.12 | −1,172.60 | −1,154.80 | −1,153.77 | **−1,150.58** |
| 3_DivMig_8_Sim | −11,928.42 | −11,766.96 | −11,844.63 | −11,726.69 | −11,715.04 | **−11,652.18** |
| Mean cost on all datasets | −2,168.36 | −2,112.62 | −2,099.81 | **−2,084.72** | −2,188.06 | −2,144.00 |

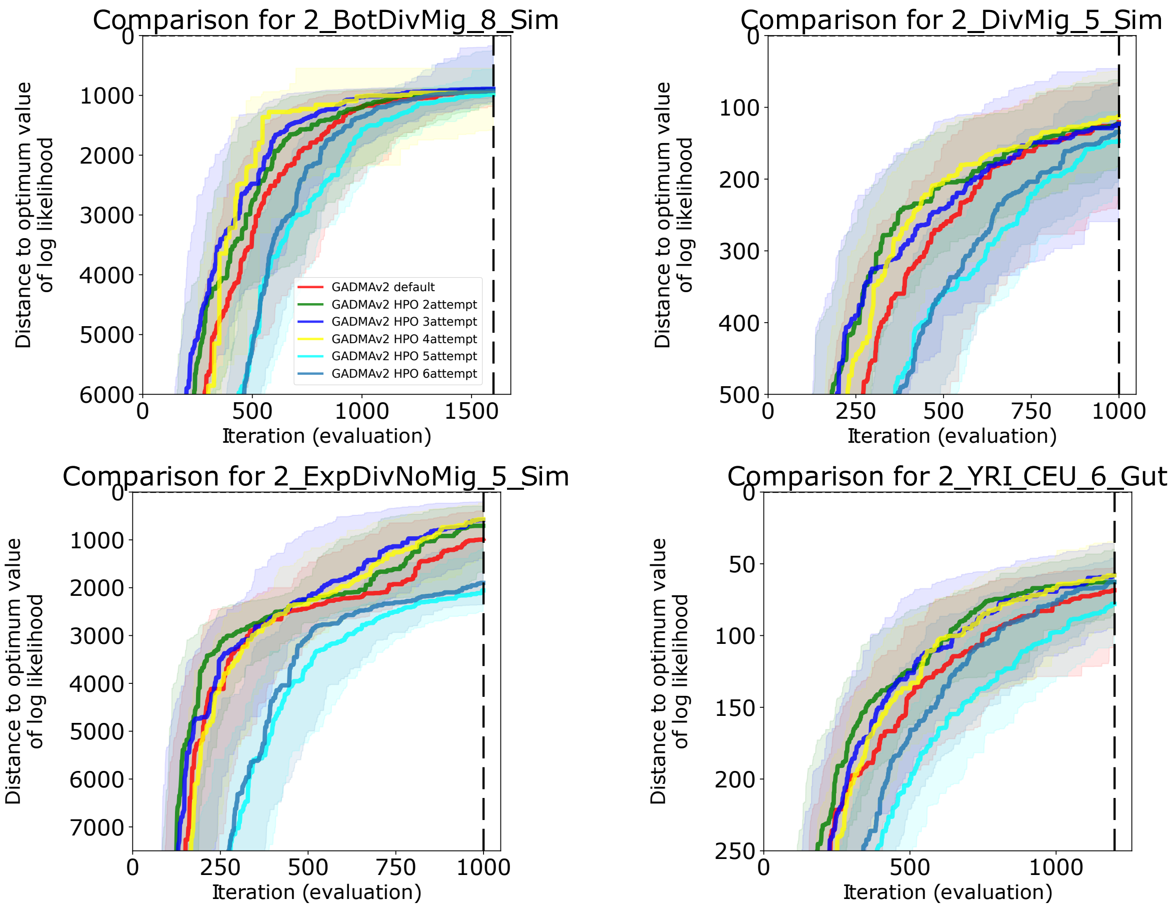

Figure S6: Convergence plots for six genetic algorithm configurations using ∂a∂i engine on four training datasets: 1) the default genetic algorithm from the initial version of GADMA, 2)-6) configurations obtained during attempts 2-6 of hyperparameter optimization with SMAC. The abscissa presents the log-likelihood evaluation number, the ordinate refers to the distance to the optimal value of log-likelihood. Solid lines correspond to median convergence over 128 runs and shadowed areas are ranges between first (0.25) and third (0.75) quartiles. The vertical dashed black line refers to the number of evaluations used to stop a genetic algorithm in SMAC.

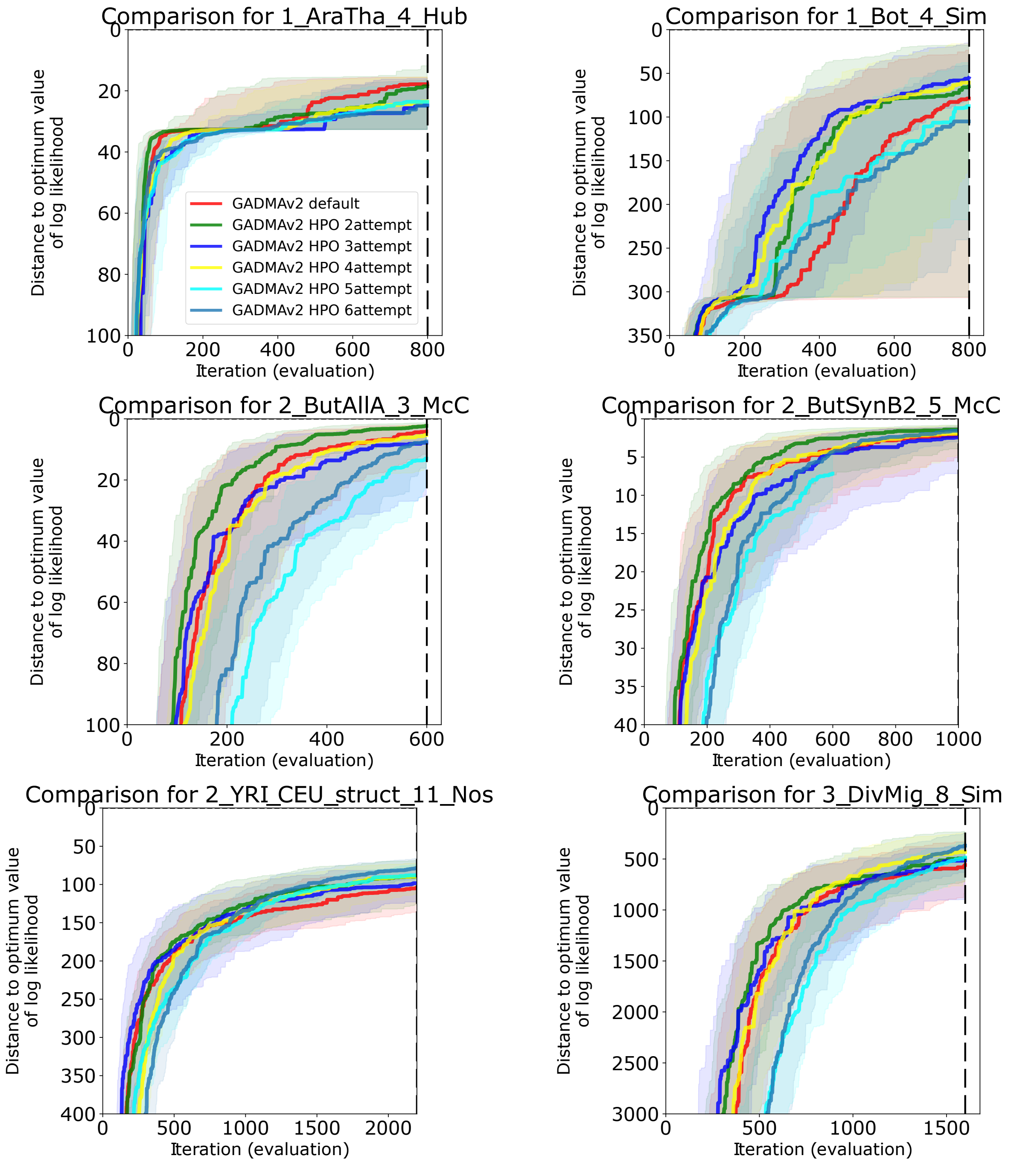

Figure S7: Convergence plots for six genetic algorithm configurations on six test datasets: 1) the default genetic algorithm from the initial version of GADMA (red colour), 2)-6) configurations obtained during attempts 2-6 of hyperparameter optimization with SMAC.

#### S1.5 SMAC costs and convergence plots using *momi2* engine

Table S4: Mean log-likelihood values (128 runs) for final configurations of six SMAC attempts on training and test datasets using *momi2* simulation engine. Genetic algorithm was stopped at the same number of evaluations used in SMAC. Two test datasets (`2_ButAllA_3_McC`, `2_ButSynB2_5_McC`) were excluded as they lacked sequence length required for *momi2* engine. Moreover, *momi2* engine does not support continuous migrations and size of ancestral population could not be inferred implicitly as for ∂a∂i and *moments*. Thus, number of parameters in datasets for *momi2* differs from *moments* and ∂a∂i. Best mean values are marked bold.

| Dataset | Par. num. | attempt number 1 (default) | 2 | 3 | 4 | 5 | 6 |
|---|---|---|---|---|---|---|---|
| Mean cost on training datasets | | −485,627.85 | −485,343.30 | **−485,225.21** | −485,432.47 | −485,728.26 | −485,515.65 |
| training datasets: | | | | | | | |
| `2_BotDivMig_8_Sim` | 7 | −363,769.82 | −363,480.25 | **−362,972.57** | −363,383.18 | −363,685.63 | −363,590.86 |
| `2_DivMig_5_Sim` | 4 | −352,307.58 | **−352,137.50** | −352,359.24 | −352,267.29 | −352,412.27 | −352,406.86 |
| `2_ExpDivNoMig_5_Sim` | 6 | −1,162,516.73 | −1,161,866.86 | **−1,161,679.79** | −1,162,153.62 | −1,162,910.47 | −1,162,178.52 |
| `2_YRI_CEU_6_Gut` | 6 | −63,917.28 | **−63,866.37** | −63,889.26 | −63,925.77 | −63,904.66 | −63,886.36 |
| Mean cost on test datasets | | −261,011.47 | −260,960.01 | −260,883.68 | −260,957.31 | −260,914.32 | **−260,856.28** |
| Test datasets: | | | | | | | |
| `1_Bot_4_Sim` | 5 | −109,574.32 | **−109,554.94** | −109,558.06 | −109,567.70 | −109,615.11 | −109,593.22 |
| `1_AraTha_4_Hub` | 5 | −228,328.17 | −228,320.02 | −228,318.47 | −228,335.39 | −228,325.95 | **−228,315.27** |
| `2_YRI_CEU_str_11_Nos` | 10 | −63,904.25 | −63,893.87 | −63,891.60 | **−63,878.87** | −63,885.36 | −63,881.22 |
| `3_DivMig_8_Sim` | 6 | −642,239.14 | −642,071.21 | −641,766.61 | −642,047.26 | −641,826.97 | **−641,635.42** |
| Mean cost on all datasets | | −350,858.02 | −350,713.32 | **−350,620.30** | −350,747.37 | −350,839.90 | −350,720.03 |

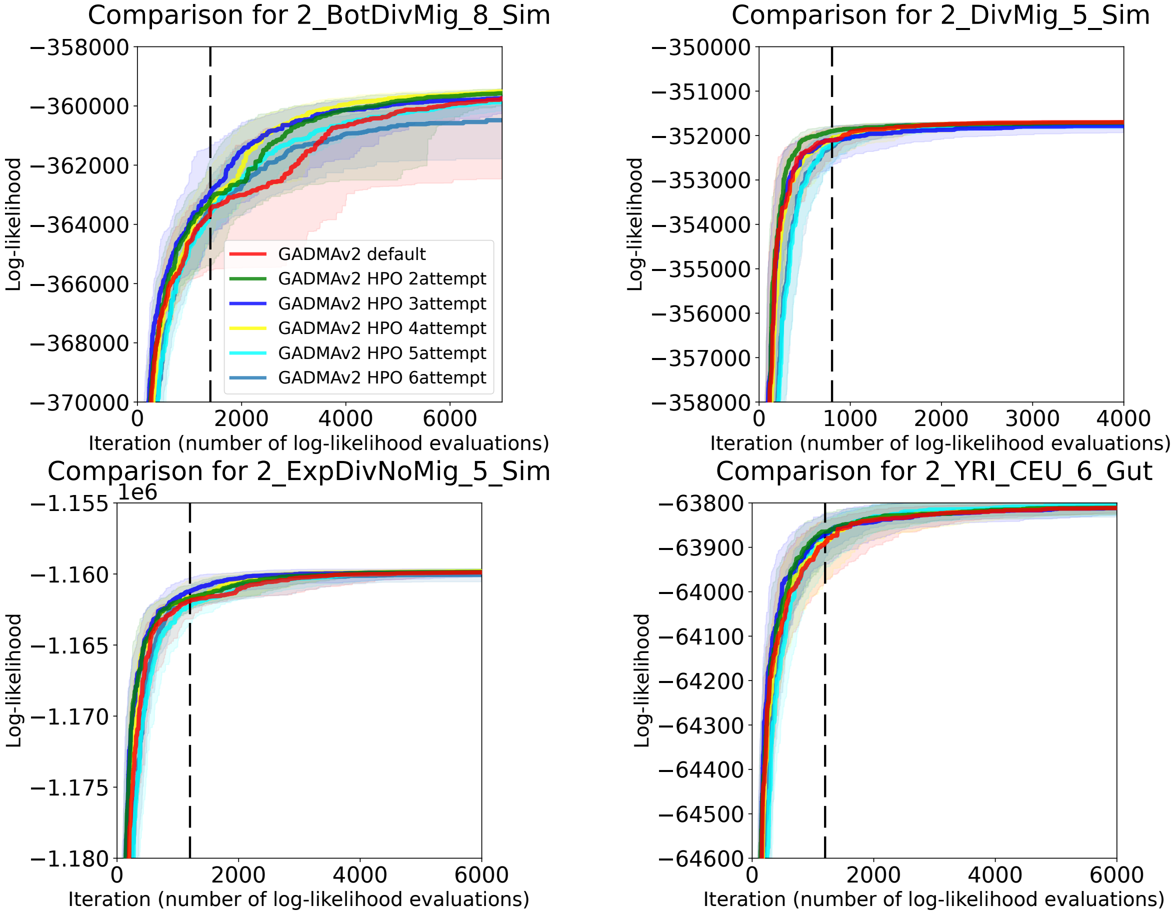

Figure S8: Convergence plots for six genetic algorithm configurations using *momi2* engine on four training datasets: 1) the default genetic algorithm from the initial version of GADMA (red colour), 2)-6) configurations obtained during attempts 2-6 of hyperparameter optimization with SMAC. The abscissa presents the log-likelihood evaluation number, the ordinate refers to the distance to the optimal value of log-likelihood. Solid lines correspond to median convergence over 128 runs and shadowed areas are ranges between first (0.25) and third (0.75) quartiles. The vertical dashed black line refers to the number of evaluations used to stop a genetic algorithm in SMAC.

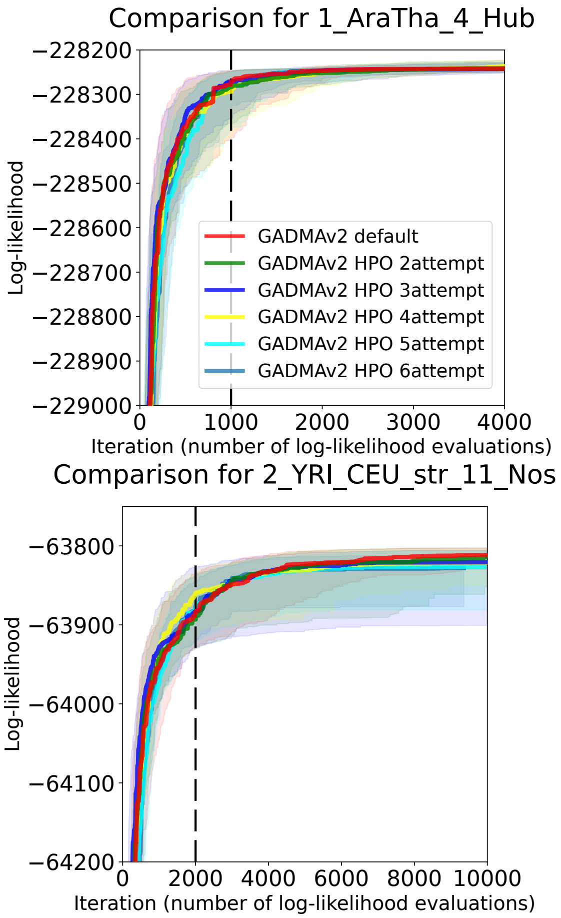

Figure S9: Convergence plots for six genetic algorithm configurations on four test datasets: 1) the default genetic algorithm from the initial version of GADMA (red colour), 2)-6) configurations obtained during attempts 2-6 of hyperparameter optimization with SMAC. Two datasets (`2_ButAllA_3_McC`, `2_ButSynB2_5_McC`) were excluded as they are not supported by *momi2* engine.

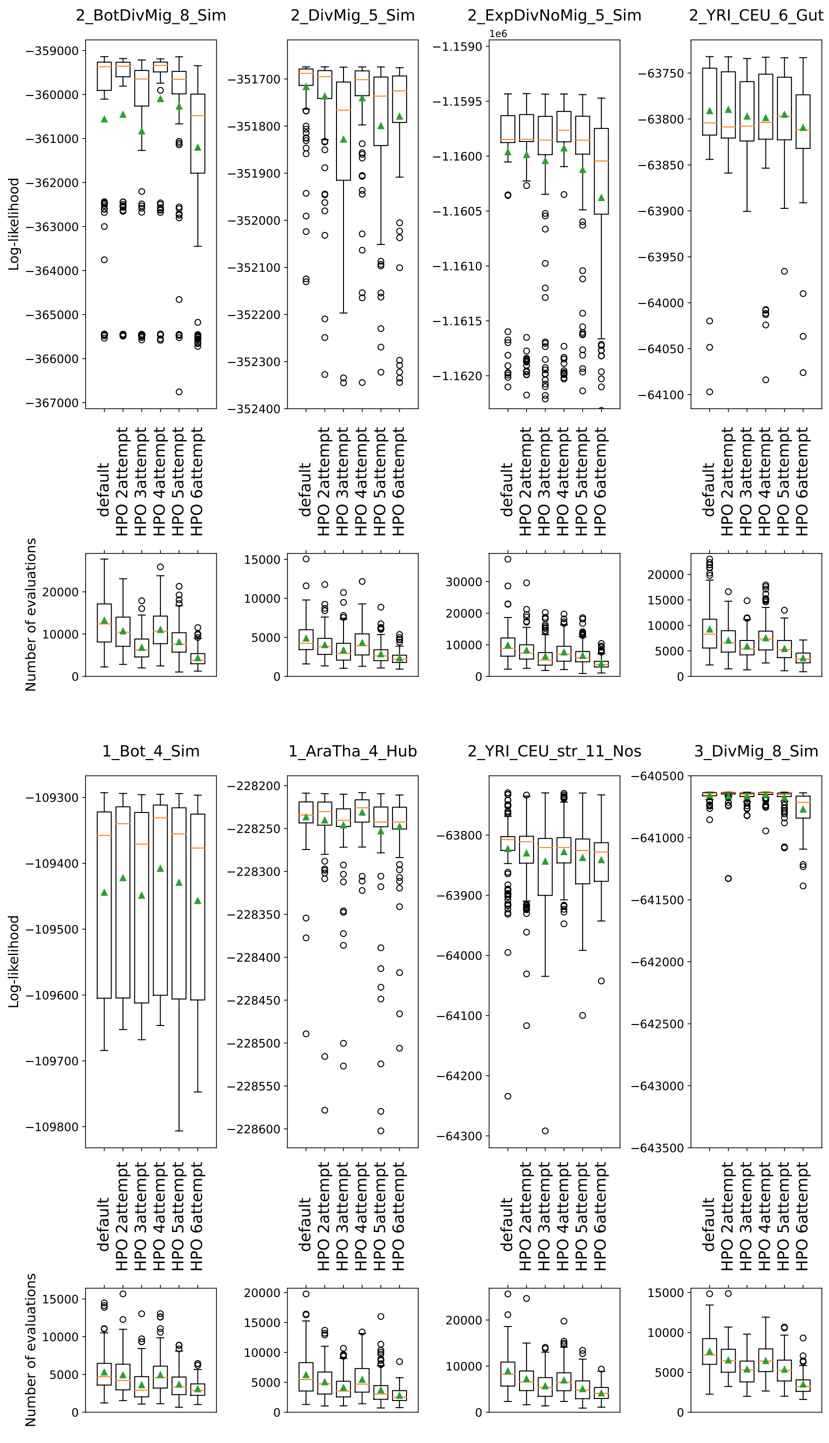

Figure S10: Boxplots of the eventual log-likelihoods and number of evaluations required for full runs of genetic algorithm with six configurations using *momi2* engine. For each dataset two plots are presented: 1) the top plot shows distribution of 128 resulting log-likelihood values; 2) the bottom plot corresponds to the distribution of the evaluations' number required for genetic algorithms to terminate. Orange line on boxplot refers to the median value, greed triangle demonstrates the mean value.

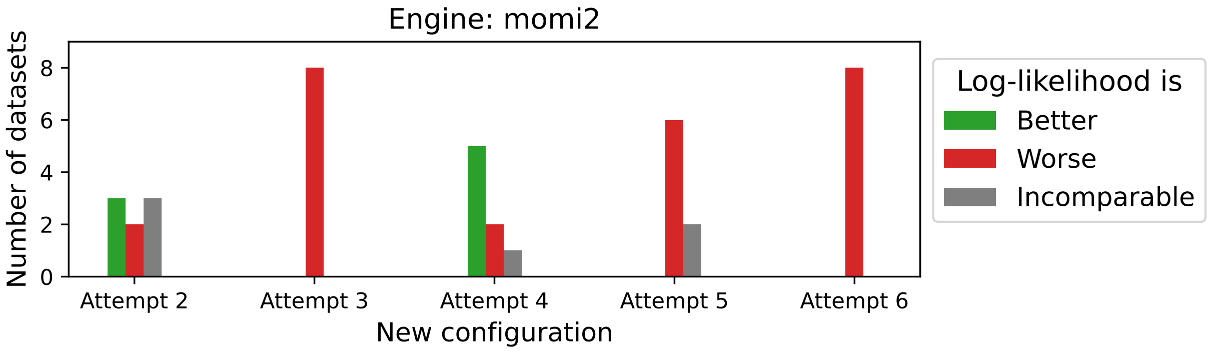

Figure S11: For each dataset, the performance of the new configurations from attempts 2-6 is categorized as either better, worse, or incomparable to the performance of the default configuration when using the *momi2* engine. The histogram illustrates the count of datasets falling into each category for each configuration. Performance is considered to be better if both the median value and both quartiles of log-likelihoods are higher than those of the default configuration. Conversely, if both the median and quartiles are lower, the dataset is categorized as worse. Otherwise, the comparison is considered to be similar or undefined.

### S2 Performance comparison of GADMA2 engines

#### S2.1 Fruit fly simulated dataset

Generally, all engines infer demographic parameters close to the ground truth values for the DROS-NOMIG model (Table 2). Engine *momi2* demonstrates the best estimations among tested engines, especially for the size of the ancestral population $N_{anc}$ and two time parameters: $T_{split}$ and $T_{AFR}$. Other engines approximate $N_{anc}$ to be lower than ground truth value of $1,720,600$ individuals: ∼1,580,000 for *moments* and ∂a∂i and ∼1,200,000 for *momentsLD*. Estimates for time parameters are also similar between ∂a∂i and *moments* but differ for other engines. European bottleneck is approximated by ∂a∂i and *moments* to start earlier 160,000 − 170,000 generations ago and end lately ∼135,000 generations ago compared to the ground truth interval from 158,000 to 154,600 generations ago. Engines *momi2* and *momentsLD* predict bottleneck time closer to the ground truth. The European population's size during the bottleneck equal to 2,200 in simulation is approximated relatively accurately by the *momentsLD* engine. Engines ∂a∂i and *moments* provide a value of ∼20,000 individuals, *momi2* engine estimate it to be lower: ∼500 individuals.

Including continuous migrations for the DROS-MIG model falls into a more complex optimization problem. As a result, the best model parameters obtained within eight runs for each tested engine have a worse log-likelihood value than the best scenarios for the DROS-NOMIG model (Table S6). However, final histories can catch the general history similar to the DROS-NOMIG model and indicate zero or low migration rates. Thus, based on these results, it is possible to build a hypothesis of population isolation and test further models without migration.

All engines provide estimations of ancestral size for DROS-STRUCT-NOMIG models similar to the estimations for models DROS-NOMIG and DROS-MIG (Table S7). Regarding population size dynamics, all engines inferred sudden expansion of the ancestral population and constant size of the African population after the split. The history of the European population is approximated by constant size using ∂a∂i and by exponential dynamic using other engines. Engines *moments* and *momentsLD* provide good approximations of the ground truth history (Figure S12). The African population history in the simulation is a sudden expansion from 1,720,600 to 8,603,000. Both engines approximate size after expansion slightly to be lower (5,000,000 − 6,500,000). The time of the African population after the split is estimated correctly by *momentsLD* and overestimated (13,000,000) by *moments*. Result histories for *moments* and *momentsLD* differ in the rate of exponential dynamic approximation for the European population history. We perform a comparison of those approximations and ground truth two-epoch history by several metrics in order to validate estimations. This comparison is presented in section S2.1.1.

During our further analysis, it turns out that all AFS-based likelihood engines, ∂a∂i, *moments* and *momi2*, demonstrate very rough common local optimums for the model DROS-STRUCT-NOMIG. Although *moments*' result is not an optimal solution, it is close to the global optimum and provides a good approximation compared to other AFS-based engines. More details about local and global optimums are available in section S2.1.2. The alternative history from local optimum of ∂a∂i and *momi2* demonstrates only slight increase in ancestral population size from 1,500,000 up to 1,800,000 - 2,000,000 individuals. The size of the African population after the divergence is estimated to be so big that it hits the upper bound. Nevertheless, the history of the European population is approximated relatively accurately. Engine ∂a∂i shows the size of the European population as a constant size of $\sim 200,000$.

The migration included in model DROS-STRUCT-MIG additionally hinders the optimization.

Engine *moments* could not find the same local optimum as for the DROS-STRUCT-NOMIG model and was stuck in a worse local optimum similar to ∂a∂i and *momi2*. The ground truth history was reconstructed the best by *momentsLD*. Despite the local optimum, all histories demonstrate low or close to zero estimations of migration rates. The sizes of the ancestral population correspond to the estimations from other analyzed models.

##### S2.1.1 Comparison of the European population histories inferred by *moments* and *momentsLD* within the DROS-STRUCT-NOMIG model

Engines *moments* and *momentsLD* provide result histories for the DROS-STRUCT-NOMIG model with parameters close to the ground truth values. However, two-epoch history of the European population after split is approximated by the exponential growth with a rate that differs between engines. In order to validate the accuracy of predicted approximations we use several simple metrics that are reported in Table S7. The notation of parameters that are used further is also available in Table S7.

First, we compute $A^{EUR}$ — mean size of European population after split, which is equal:

$$A^{EUR} = \begin{cases} \dfrac{N_{EUR0} \cdot (T_{split} - T_{EUR}) + N_{EUR} \cdot T_{EUR}}{T_{split}}, & \text{for ground truth two-epoch history,} \\ \dfrac{N_{EUR0}(\exp\{-r \cdot T_{split}\} - 1)}{-r \cdot T_{split}}, & \text{for exponential growth history,} \end{cases}$$

where $r = -\frac{\log N_{EUR} - \log N_{EUR0}}{T_{split}}$.

However, mean population size differs a lot between tested histories: 1,051,914 individuals for ground truth history, 391,566 for the history from *moments* and 13,041,907 for the history from *momentsLD*. Based on received estimations the history from *moments* has the average size of the European population closer to the ground truth than the history from *momentsLD*.

We note that such a metric as mean size $A^{EUR}$ almost do not reflect the severe bottleneck of the ground truth history of the European population after split. Thus, we evaluate another metric — the harmonic mean, that is often used to approximate the overall effective population size from the recent fluctuations. In general, it sums the inverse population sizes and gives more weight to small values. The harmonic mean can be easily extended by using integrals when size of population is a continuous function such as exponential growth. To evaluate the harmonic mean size $H^{EUR}$ of the European population in our histories we use the following:

$$\frac{1}{H^{EUR}} = \begin{cases} \dfrac{1}{T_{split}} \left[ \dfrac{T_{split} - T_{EUR}}{N_{EUR0}} + \dfrac{T_{EUR}}{N_{EUR}} \right], & \text{for ground truth two-epoch history,} \\ \dfrac{1}{T_{split}} \left[ \dfrac{(\exp\{r \cdot T_{split}\} - 1)}{N_{EUR0} \cdot r} \right], & \text{for exponential growth history,} \end{cases}$$

where $r = -\frac{\log N_{EUR} - \log N_{EUR0}}{T_{split}}$.

Harmonic mean of the exponential growth history obtained for *momentsLD* engine (93,617) is very close to the harmonic mean size from the ground truth history (93,531). History received for *moments* engine has greater (132,932) harmonic mean size than other tested scenarios.

Finally, we estimate and compare histories by the mean time of coalescence. Using *msprime* we simulate 100,000 coalescent trees under each history and then report the average tree height. This characteristic is also referred to as the time of the most recent common ancestor and is denoted as $t^{EUP}_{MRCA}$. For the simulations we use the same number of samples as in our data which is 10 chromosomes. In contrast to the two previous metrics, time of coalescence is estimated for the entire history of the European population including the history of the ancestral population of European and African populations. However, we ensure that $t^{EUR}_{MRCA}$ is mostly formed by recent history of European population after split by fixing the history of the ancestral population to ground truth and obtaining similar estimations. Based on the results *moments* engine has a mean coalescent time of 330,932 generations which is closer to the ground truth estimation of 1,075,080 than the 193,677 generations obtained for the *momentsLD* history. However, we note that coalescent times are quite different for the ground truth two-epoch history and for exponential approximations.

Table S5: Models of *Drosophila melanogaster* populations' history and GADMA2's likelihood engines used for performance comparison. If engine was used to infer model parameters then notation "+" is set between these engine and model, otherwise notation "−" is specified. Engine *momi2* is not compared for the models with migration as it does not support continuous migrations.

| Model | ∂a∂i | *moments* | *momi2* | *momentsLD* |
|---|---|---|---|---|
| DROS-NOMIG | + | + | + | + |
| DROS-MIG | + | + | − | + |
| DROS-STRUCT-NOMIG | + | + | + | + |
| DROS-STRUCT-MIG | + | + | − | + |

Table S6: The demographic parameters of *Drosophila melanogaster* history with migration (DROS-MIG model) inferred with different engines in GADMA2. Ground truth are the parameter values from the original paper Li and Stephan (2006) used in simulation powered by *stdpopsim* (Adrion et al., 2020). Engine *momi2* is excluded as it does not support continuous migrations. Log-likelihood values are not comparable between different engines.

| | Ground truth | ∂a∂i | *moments* | *momentsLD* |
|---|---|---|---|---|
| Log-likelihood: | | −3,552.69 | −2,047.45 | −576.37 |
| Parameters: | | | | |
| $N_{anc}$ | 1,720,600 | 1,595,786 | 1,588,257 | 1,269,211 |
| $N_{AFR}$ | 8,603,000 | 8,349,726 | 8,843,033 | 8,033,141 |
| $N_{EUR0}$ | 2,200 | 70,023 | 45,297 | 6,891 |
| $N_{EUR}$ | 1,075,000 | 1,342,486 | 1,588,257 | 1,368,298 |
| $m_{AFR-EUR}$ | 0 | 0 | 0 | 7*10-13 |
| $m_{EUR-AFR}$ | 0 | 6*10-8 | 0 | 5*10-9 |
| $T_{AFR}$ | 600,000 | 305,417 | 529,111 | 715,581 |
| $T_{split}$ | 158,000 | 115,838 | 188,875 | 153,346 |
| $T_{EUR}$ | 154,600 | 56,331 | 112,448 | 142,100 |

$N_{anc}$: size of ancestral population; $N_{AFR}$: size of African population after expansion; $N_{EUR0}$: European bottleneck population size after divergence; $N_{EUR}$: modern size of European population; $m_{AFR-EUR}$: migration rate from African population to European population; $m_{EUR-AFR}$: migration rate from European population to African population; $T_{AFR}$: time of African size expansion in generations; $T_{split}$: time of divergence in generations; $T_{EUR}$: time of European expansion in generations.

Table S7: The demographic parameters of *Drosophila melanogaster* history without migration for structure (2, 1) (DROS-STRUCT-NOMIG model) inferred with different engines in GADMA2. Ground truth are the parameter values from the original paper Li and Stephan (2006) used in simulation powered by *stdpopsim* (Adrion et al., 2020). Log-likelihood values are not comparable between different engines.

| | Ground truth | ∂a∂i | *moments* | *momi2* | *momentsLD* |
|---|---|---|---|---|---|
| Log-likelihood: | | −17,321.62 | −5,290.52 | −53,500,374.42 | −1,249.91 |
| Parameters: | | | | | |
| $N_{anc}$ | 1,720,600 | 1,561,905 | 1,583,371 | 1,726,753 | 1,229,777 |
| $N_{anc\_exp}$ | 8,603,000 | 1,908,519 | 5,098,605 | 2,643,656 | 6,554,260 |
| $N_{AFR}$ | 8,603,000 | 156,190,553* | 12,847,325 | 172,667,794* | 8,839,943 |
| $N_{EUR0}$ | 2,200† | 195,933 | 33,988 | 46,965 | 9,934 |
| $N_{EUR}$ | 1,075,000† | 195,933 | 1,522,854$^{exp}$ | 1,161,462$^{exp}$ | 122,906,664$^{*,exp}$ |
| $T_{AFR}$ | 600,000 | 1,720,902 | 574,107 | 976,673 | 756,507 |
| $T_{split}$ | 158,000 | 354,058 | 234,750 | 280,951 | 155,714 |
| Characteristics: | | | | | |
| $A^{EUR}$ | 1,051,914 | | 391,566 | | 13,041,907 |
| $H^{EUR}$ | 93,531 | | 132,183 | | 93,617 |
| $t^{EUR}_{MRCA}$ | 1,075,080 | | 330,932 | | 193,677 |

$N_{anc}$: size of ancestral population; $N_{anc\_exp}$: size of ancestral population after expansion; $N_{AFR}$: size of African population after divergence; $N_{EUR0}$: European population size after divergence; $N_{EUR}$: modern European population size; $T_{AFR}$: time of ancestral population size expansion in generations; $T_{split}$: time of divergence in generations.
†Size of European population after divergence was $N_{EUR0}$ for 3,400 generations and then expanded up to $N_{EUR}$.
*Estimation hits the upper bound of the parameter.
$^{exp}$Exponential growth.
$A^{EUR}$ — arithmetic mean size of European population after split.
$H^{EUR}$ — harmonic mean size of European population after split.
$t^{EUR}_{MRCA}$ — mean time in generations of the most recent common ancestor estimated from 100,000 simulated coalescent trees for the full history of European population.

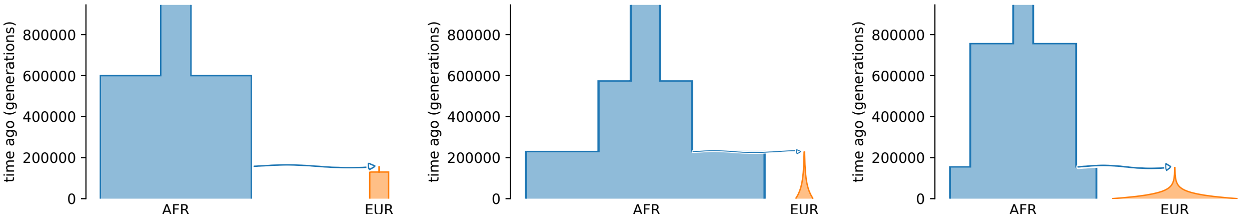

(a) Ground truth demographic history used in simulation (b) Demographic history inferred by *moments* engine (c) Demographic history inferred by *momentsLD* engine

Figure S12: Comparison of the original demographic history of *Drosophila melanogaster* and approximations for DROS-STRUCT-NOMIG model by *moments* and *momentsLD* engines.

Table S8: The demographic parameters of *Drosophila melanogaster* history with migration for structure (2, 1) (DROS-STRUCT-MIG model) inferred with different engines in GADMA2. Ground truth are the parameter values from the original paper Li and Stephan (2006) used in simulation powered by *stdpopsim* (Adrion et al., 2020). Engine *momi2* is excluded as it does not support continuous migrations. Log-likelihood values are not comparable between different engines.

| | Ground truth | ∂a∂i | *moments* | *momentsLD* |
|---|---|---|---|---|
| Log-likelihood: | | −16,934.97 | −18,576.64 | −1,743.76 |
| Parameters: | | | | |
| $N_{anc}$ | 1,720,600 | 1,561,773 | 1,562,147 | 1,438,130 |
| $N_{anc\_exp}$ | 8,603,000 | 1,871,611 | 2,119,711$^{lin}$ | 6,644,707 |
| $N_{AFR}$ | 8,603,000 | 156,177,325$^{*}$ | 156,214,765$^{*}$ | 10,088,036 |
| $N_{EUR0}$ | 2,200$^{\dagger}$ | 185,084 | 67,323 | 23,528 |
| $N_{EUR}$ | 1,075,000$^{\dagger}$ | 204,561$^{exp}$ | 664,550$^{exp}$ | 3,126,005$^{exp}$ |
| $m_{AFR-EUR}$ | 0 | $2 \cdot 10^{-8}$ | $7 \cdot 10^{-9}$ | $8 \cdot 10^{-9}$ |
| $m_{EUR-AFR}$ | 0 | $5 \cdot 10^{-11}$ | $3 \cdot 10^{-10}$ | ∼0 |
| $T_{AFR}$ | 600,000 | 1,880,793 | 1,768,872 | 1,244,572 |
| $T_{split}$ | 158,000 | 357,484 | 309,256 | 178,351 |

$N_{anc}$: size of ancestral population; $N_{anc\_exp}$: size of ancestral population after expansion; $N_{AFR}$: size of African population after divergence; $N_{EUR0}$: European population size after divergence; $N_{EUR}$: modern European population size; $m_{AFR-EUR}$: migration rate from African population to European population; $m_{EUR-AFR}$: migration rate from European population to African population; $T_{AFR}$: time of ancestral population size expansion in generations; $T_{split}$: time of divergence in generations.

$^{\dagger}$Size of European population after divergence was $N_{EUR0}$ for 3,400 generations and then expanded up to $N_{EUR}$.

$^{*}$Estimation hits the upper bound of the parameter.

$^{exp}$Exponential growth.

$^{lin}$Linear growth.

Table S9: Comparison of average times for log-likelihood evaluation between used GADMA2's likelihood engines and models of *Drosophila melanogaster* history. For each model and likelihood engine mean value and standard deviation are reported.

| Model | ∂a∂i | *moments* | *momi2* | *momentsLD* |
|---|---|---|---|---|
| DROS-NOMIG | $1.09 \pm 22.31$ | $0.11 \pm 0.18$ | $0.01 \pm 0.006$ | $0.84 \pm 0.82$ |
| DROS-MIG | $0.68 \pm 5.96$ | $0.05 \pm 0.07$ | – | $0.80 \pm 0.79$ |
| DROS-STRUCT-NOMIG | $0.19 \pm 2.19$ | $0.08 \pm 0.13$ | $0.02 \pm 0.01$ | $11.10 \pm 33.09$ |
| DROS-STRUCT-MIG | $0.38 \pm 4.75$ | $0.27 \pm 0.49$ | – | $17.62 \pm 45.24$ |

Table S10: Mean number of log-likelihood evaluations required for demographic inference averaged over eight GADMA2 runs for each engine and model of *Drosophila melanogaster* history.

| Model | ∂a∂i | *moments* | *momi2* | *momentsLD* |
|---|---|---|---|---|
| DROS-NOMIG | 11,668 | 13,607 | 8,345 | 12,277 |
| DROS-MIG | 3,849 | 9,438 | – | 13,428 |
| DROS-STRUCT-NOMIG | 5,311 | 6,457 | 1,564 | 12,230 |
| DROS-STRUCT-MIG | 7,871 | 9,241 | – | 12,007 |

##### S2.1.2 Investigating local optimum phenomena

We observe that AFS-based engines were stuck in local optimum during inference of fruit flies demographic history using misspecified models with structure (2, 1). However, we should make sure that result histories are local optimums. In order to do it, we construct new models that align with the previous structure models but contain some additional restrictions or fixations. Therefore, obtaining parameters with better log-likelihood values for these new models will prove that previous results are not optimal. We will perform our analysis for the models without migrations. We believe the same results could be demonstrated for the model that includes migrations.

We take the model DROS-STRUCT-NOMIG as a basic model. Using GADMA2's API, we construct two new models: DROS-STRUCT-NOMIG-AFR and DROS-STRUCT-NOMIG-AFR-ANC. The first model DROS-STRUCT-NOMIG-AFR is the model with structure (2, 1) but obliges the size of the ancestral population before the split to be equal to the size of the African population that is constant after the split. The second model DROS-STRUCT-NOMIG-AFR-ANC is the first model with one more fixation: ancestral population experiences sudden growth. The latter means that the dynamic during the second epoch of the ancestral population is fixed to a constant size.

Parameters inferred for the DROS-STRUCT-NOMIG-AFR model are alike for all three AFS-based engines (Table S11). Estimations are very similar between *moments* and *momi2* and differ a little for ∂a∂i. As it turns out further, the ∂a∂i engine could not reach global optimum again. Population size dynamics were inferred the same for all tested engines and are the following: ancestral population before the split and Europen population after the split experience exponential growth. We note that the dynamic of the African population was fixed to constant size in the analyzed model. The size of African population is estimated to be higher: ∼10,000,000 vs 8,603,000 from simulation. European population history is approximated by exponential growth from 33,000 to 1,500,000 for ∂a∂i engine and from 17,000 − 20,000 to 6,000,000 for *moments* and *momi2* engines. The time of ancestral exponential growth is estimated earlier ∼750,000 than the ground truth value of 600,000. Value for the population split time is close to the ground truth for *moments* and *momi2*.

The last DROS-STRUCT-NOMIG-AFR-ANC model has a dynamic for ancestral population expansion as sudden change with constant size. The result parameters are very consisted between AFS-based engines (Table S12). The ancestral size and size of the African population are estimated to be very close to the actual value. European population history is an exponential growth from ∼17,000 up to ∼ 7,000,000 individuals. Estimations for the time of ancestral population instantaneous expansion and the time of split are accurate for ∂a∂i and *momi2*. However, *moments* predict them to be rather late.

Finally, we check for the result log-likelihood values to understand which history is an actual global optimum for each AFS-based engine. In the case of ∂a∂i engine, the best history with a log-likelihood value of −3,541 is obtained for the last DROS-STRUCT-NOMIG-AFR-ANC model. In the case of DROS-STRUCT-NOMIG and DROS-STRUCT-NOMIG-AFR models, the best achieved by ∂a∂i log-likelihood values are −17,321 and −7,328 correspondingly. For *moments* and *momi2*, the best result is for the DROS-STRUCT-NOMIG-AFR model. In the case of *moments* best achieved log-likelihood value is 1,409 compared to values of −5,290 and −1,800 for models DROS-STRUCT-NOMIG and DROS-STRUCT-NOMIG-AFR-ANC respectively. Engine *momi2* provide log-likelihoods: −53,490,284 for DROS-STRUCT-NOMIG-AFR model, −53,490,608 for DROS-STRUCT-NOMIG-AFR-ANC model and −53,500,374 for DROS-STRUCT-NOMIG model.

Thus, we proved that AFS-based engines failed to reach the existing global optimum within eight runs for the DROS-STRUCT-NOMIG model.

Table S11: The demographic parameters of *Drosophila melanogaster* history without migration for structure (2, 1) and restriction on population sizes of ancestral and African populations (DROS-STRUCT-NOMIG-AFR model) inferred with different AFS-based engines in GADMA2. Ground truth are the parameter values from the original paper Li and Stephan (2006) used in simulation powered by *stdpopsim* (Adrion et al., 2020). Log-likelihood values are not comparable between different engines.

| | Ground truth | ∂a∂i | *moments* | *momi2* |
|---|---|---|---|---|
| Log-likelihood: | | −7,328 | −1,409 | −53,490,284 |
| Parameters: | | | | |
| $N_{anc}$ | 1,720,600 | 1,603,921 | 1,577,475 | 1,722,290 |
| $N_{anc2}$ | 8,603,000 | 10,707,776$^{exp}$ | 10,018,543$^{exp}$ | 11,090,692$^{exp}$ |
| $N_{AFR}$ (=$N_{anc2}$) | 8,603,000 | 10,707,776 | 10,018,543 | 11,090,692 |
| $N_{EUR0}$ | 2,200$^{\dagger}$ | 33,682 | 17,352 | 19,472 |
| $N_{EUR}$ | 1,075,000$^{\dagger}$ | 1,464,379$^{exp}$ | 5,876,094$^{exp}$ | 6,171,224$^{exp}$ |
| $T_{AFR}$ | 600,000 | 721,764 | 738,258 | 774,321 |
| $T_{split}$ | 158,000 | 234,172 | 182,987 | 197,924 |

$N_{anc}$: size of ancestral population; $N_{anc2}$: size of ancestral population after expansion; $N_{AFR}$: size of African population after divergence; $N_{EUR0}$: European population size after divergence; $N_{EUR}$: modern European population size; $T_{AFR}$: time of ancestral population size expansion in generations; $T_{split}$: time of divergence in generations.
$^{\dagger}$Size of European population after divergence was $N_{EUR0}$ for 3,400 generations and then expanded up to $N_{EUR}$.
$^{exp}$Exponential growth.

Table S12: The demographic parameters of *Drosophila melanogaster* history without migration for structure (2, 1) and restrictions on instantaneous expansion of ancestral population and on population sizes of ancestral and African populations (DROS-STRUCT-NOMIG-AFR-ANC model) inferred with different AFS-based engines in GADMA2. Ground truth are the parameter values from the original paper Li and Stephan (2006) used in simulation powered by *stdpopsim* (Adrion et al., 2020). Log-likelihood values are not comparable between different engines.

| | Ground truth | ∂a∂i | *moments* | *momi2* |
|---|---|---|---|---|
| Log-likelihood: | | −3,541 | −1,800 | −53,490,608 |
| Parameters: | | | | |
| $N_{anc}$ | 1,720,600 | 1,599,932 | 1,583,846 | 1,727,916 |
| $N_{AFR}$ | 8,603,000 | 8,370,165 | 8,275,637 | 9,191,357 |
| $N_{EUR0}$ | 2,200$^{\dagger}$ | 16,888 | 16,459 | 17,967 |
| $N_{EUR}$ | 1,075,000$^{\dagger}$ | 6,720,978$^{exp}$ | 7,320,777$^{exp}$ | 7,607,237$^{exp}$ |
| $T_{AFR}$ | 600,000 | 600,901 | 446,265 | 574,998 |
| $T_{split}$ | 158,000 | 181,294 | 76,707 | 191,246 |

$N_{anc}$: size of ancestral population; $N_{AFR}$: size of ancestral population after instantaneous growth and size of African population after divergence; $N_{EUR0}$: European population size after divergence; $N_{EUR}$: modern European population size; $T_{AFR}$: time of ancestral population size expansion in generations; $T_{split}$: time of divergence in generations.
$^{\dagger}$Size of European population after divergence was $N_{EUR0}$ for 3,400 generations and then expanded up to $N_{EUR}$.
$^{exp}$Exponential growth.

#### S2.2 Orangutan simulated dataset

Engines provide similar histories for a misspecified ORAN-NOMIG model lacking migrations (Table S14). The results for AFS-based engines and the *momentsLD* engine differ slightly for ancestral population size and size of Sumatran orangutan after the split. The ancestral population size with a ground truth value of 17,934 individuals is estimated to be ∼19,500 by ∂a∂i, *moments* and *momi2*. Engine *momentsLD* provides a value of ∼14,500 that is lower than the value used for simulation. AFS-based engines approximate the size of Sumatran orangutans after divergence as ∼8,000 individuals, *momentsLD* provide a value of 5,830 compared to the true value of 7,217. For other parameters, the predictions between all engines are alike. The time of divergence is approximated by the value of 12,000 generations ago compared to the 20,157 used for simulation. The modern sizes of populations are overestimated, for example, the modern size of the Sumatran population is estimated by ∼55,000 value when the ground truth is 37,661 individuals. All engines underestimate the Bornean population size after the split by the value of ∼6,500 compared to the 10,617 used in simulations. Based on the inferred sized for Bornean populations, all engines predict slow exponential growth of the Bornean population from ∼6,500 up to ∼10,500. However, the original history of the Bornean population contains a slight exponential decline from 10,617 to 8,805.

In the case of the ORAN-MIG model that aligns with ground truth history, all engines provide close to truth parameters (Table S15). Log-likelihood values are better than for model ORAN-NOMIG showing that this model better fits the data.

Model ORAN-STRUCT-NOMIG with structure (1, 1) allows automatic inference of the population size dynamics. Nevertheless, it lacks migrations like the ORAN-NOMIG model. The result demographic histories are very similar between engines and the result of the ORAN-NOMIG model. However, dynamics for Bornean population size are inferred constant by *moments* and *momi2* engines. Other dynamics are approximated correctly as exponential. As the originally Bornean population had a slight exponential decrease in population size, approximation by constant size is relatively good. However, the value of log-likelihood for the ORAN-STRUCT-NOMIG model using *moments* and *momi2* is not as good as the value for the ORAN-NOMIG model. Therefore, we must investigate if the results are optimal or underoptimized, like fruit flies' results.

The difference between models ORAN-NOMIG and ORAN-STRUCT-NOMIG is the following. The ORAN-STRUCT-NOMIG model obliges the sum of Bornean and Sumatran population sizes after the split to equal the ancestral population size before the split. We use this restriction as it is valid for the ground truth history. However, our first ORAN-NOMIG model does not follow this restriction, and sizes were inferred independently. From the parameters in Table S14, we can conclude that the result demographic parameters for the ORAN-NOMIG model do not satisfy this rule for any engine. In order to prove that the constant dynamic for the Bornean population inferred by *moments* and *momi2* for the ORAN-STRUCT-NOMIG model is a consequence of the model's restriction on population sizes, we perform two additional experiments with modified models. First, we checked that the result ORAN-STRUCT-NOMIG model with constant size has better value of log-likelihood than inferred history for modified ORAN-NOMIG model with restriction on population sizes. It ensure that our results for the ORAN-STRUCT-NOMIG model are not underoptimized. Second, we demonstrate that if the restriction is removed for the ORAN-STRUCT-NOMIG model the results will show exponential growth for the Bornean population. Both experiments are conducted using *moments* and *momi2* engines.

The first experiment confirms that the result history for the modified ORAN-NOMIG model with restriction on population sizes has worse value of log-likelihood than history with the constant size obtained for ORAN-STRUCT-NOMIG model both for *moments* and *momi2* engines. The best history for the modified model in case of *moments* engine has log-likelihood of −175,106 compared to −174,279 obtained for the ORAN-STRUCT-NOMIG model. The identical situation is for *momi2* engine: maximum log-likelihood value is equal to −48,546,693 for the new model and −48,545,934 obtained for ORAN-STRUCT-NOMIG model.

The result histories from the second experiment for modified ORAN-STRUCT-NOMIG model with neglected restriction on population sizes shows exponential size change of the Bornean population and better value of log-likelihood than the original ORAN-STRUCT-NOMIG model for *moments* and *momi2* engines. In case of *momi2* engine best history for modified model has parameters close to parameters received for the ORAN-NOMIG model and presented in Table S14. Inferred dynamics for both populations are estimated correctly as exponential size changes. The log-likelihood of this history is equal to −48,541,478 which is better than −48,545,934 obtained for the original ORAN-STRUCT-NOMIG model (Table S16).

However, in the case of *moments* engine best history with a log-likelihood of −169,381 shows unexpected linear size change for the Sumatran population. We note that other parameters are close to the estimates for ORAN-NOMIG model from Table S14 and that inferred linear change (from 4,500 to 45,200 individuals) is very similar to the ground truth exponential growth (from 7,317 to 37,661 individuals). For example, mean size of Sumatran population is equal to 24,850 for received linear size change and to 18,520 for ground truth exponential size change. Moreover, the next to the best history with log-likelihood of −169,924 has exponential dynamics for both populations and other parameters close to the results for ORAN-NOMIG model (Table S14). Both those histories have log-likelihood better than the result −174,279 for the original ORAN-STRUCT-NOMIG model with the restriction on population sizes (Table S16). Using GADMA2 new feature we additionally exclude linear size change from dynamics available for the inference using *moments* and obtain correct exponential dynamics for both populations. The result parameters are also close to the estimations for ORAN-NOMIG model. All results of additional experiments using *moments* are presented in the Table S17.

Moreover, we obtain that ∂a∂i and *momentsLD* engines are consistent with *moments* and also prefer the model with linear size change for the Sumatran population over the history with exponential size change in absence of migration. Histories within a model with linear size change of Sumatran population are inferred for those engines and their log-likelihoods are compared with results for the ORAN-NOMIG model. Result parameters and best log-likelihoods are presented in Table S18. The best history with linear dynamic has better log-likelihood of −172,654 than the result history obtained for ORAN-NOMIG model with log-likelihood of −173,194.31 for ∂a∂i engine. In case of *momentsLD* history with linear size change provides log-likelihood of −9,131 compared to the value of −9,556 received for the ORAN-NOMIG model. Thus, we demonstrate that model misspecifications like absence of migrations may lead to confusions between exponential and linear dynamics for *moments*, ∂a∂i and *momentsLD* engines. We have not observed such behaviour for *momi2* engine as it does not support linear size change.

Model ORAN-STRUCT-MIG aligns with ground truth history. Inferred parameters are very close to the values used in data simulation. Population size dynamics are predicted correctly.

Table S13: Models of orangutan's history and GADMA2's likelihood engines used for performance comparison. If engine was used to infer model parameters then notation "+" is set between these engine and model, otherwise notation "−" is specified. Engine *momi2* is not compared for the models with migration as it does not support continuous migrations.

| Model | ∂a∂i | *moments* | *momi2* | *momentsLD* |
|---|---|---|---|---|
| ORAN-NOMIG | + | + | + | + |
| ORAN-MIG | + | + | − | + |
| ORAN-STRUCT-NOMIG | + | + | + | + |
| ORAN-STRUCT-MIG | + | + | − | + |
| ORAN-PULSE1 | − | − | + | − |
| ORAN-PULSE3 | − | − | + | − |
| ORAN-PULSE7 | − | − | + | − |

Table S14: The demographic parameters of orangutan history without migration (ORAN-NOMIG model) inferred with different engines in GADMA2. Ground truth are the simulated parameter values that were obtained from the original paper Locke et al. (2011).

| | Ground truth | ∂a∂i | *moments* | *momi2* | *momentsLD* |
|---|---|---|---|---|---|
| Log-likelihood: | | −173,194.31 | −169,881.74 | −48,541,453.53 | −9,556.62 |
| Parameters: | | | | | |
| $N_{anc}$ | 17,934 | 19,834 | 19,776 | 19,331 | 14,595 |
| $N_{Bor\_split}$ | 10,617 | 6,753 | 6,278 | 6,187 | 6,215 |
| $N_{Sum\_split}$ | 7,317 | 8,516 | 7,753 | 7,719 | 5,830 |
| $N_{Bor}$ | 8,805 | 11,039 | 10,886 | 10,663 | 10,432 |
| $N_{Sum}$ | 37,661 | 55,733 | 56,415 | 54,184 | 61,068 |
| $T_{split}$ (gen.) | 20,157 | 12,090 | 11,458 | 11,270 | 12,361 |

$N_{anc}$: size of ancestral population; $N_{Bor\_split}$: size of *Pongo pygmaeus* at split; $N_{Sum\_split}$: size of *Pongo abelii* at split; $N_{Bor}$: size of *Pongo pygmaeus* after exponential size change; $N_{Sum}$: size of *Pongo abelii* after exponential growth; $T_{split}$: time of divergence in generations.

Table S15: The demographic parameters of orangutan history with migration (ORAN-MIG model) inferred with different engines in GADMA2. Ground truth are the simulated parameter values that were obtained from the original paper Locke et al. (2011). *Momi2* engine was excluded as it does not support continuous migrations.

| | Ground truth | ∂a∂i | *moments* | *momentsLD* |
|---|---|---|---|---|
| Log-likelihood | | −1,189 | −1,068 | −50 |
| Parameters: | | | | |
| $N_{anc}$ | 17,934 | 17,864 | 17,945 | 17,667 |
| $N_{Bor\_split}$ | 10,617 | 10,787 | 10,246 | 10,269 |
| $N_{Sum\_split}$ | 7,317 | 7,564 | 7,216 | 7,106 |
| $N_{Bor}$ | 8,805 | 9,242 | 9,036 | 8,705 |
| $N_{Sum}$ | 37,661 | 38,800 | 37,839 | 37,179 |
| $m_{Bor-Sum}(\times 10^{-5})$ | 0.66 | 0.66 | 0.66 | 0.69 |
| $m_{Sum-Bor}(\times 10^{-5})$ | 1.10 | 1.06 | 1.07 | 1.11 |
| $T_{split}$ (gen.) | 20,157 | 20,847 | 19,916 | 19,712 |

$N_{anc}$: size of ancestral population; $N_{Bor\_split}$: size of *Pongo pygmaeus* at split; $N_{Sum\_split}$: size of *Pongo abelii* at split; $N_{Bor}$: size of *Pongo pygmaeus* after exponential size change; $N_{Sum}$: size of *Pongo abelii* after exponential size change; $m_{Bor-Sum}$: migration rate from *Pongo pygmaeus* to *Pongo abelii*; $m_{Sum-Bor}$: migration rate from *Pongo abelii* to *Pongo pygmaeus*; $T_{split}$: time of divergence in generations.

Table S16: The demographic parameters of orangutan history without migration for structure (1, 1) (ORAN-STRUCT-NOMIG model) inferred with different engines in GADMA2. Ground truth are the simulated parameter values that were obtained from the original paper Locke et al. (2011). Log-likelihood values are not comparable between different engines.

| | Ground truth | ∂a∂i | *moments* | *momi2* | *momentsLD* |
|---|---|---|---|---|---|
| Log-likelihood: | | −176,605.47 | −174,279.46 | −48,545,934.48 | −9,962.32 |
| Parameters: | | | | | |
| $N_{anc}$ | 17,934 | 19,428 | 19,512 | 19,086 | 14,096 |
| $N_{Bor\_split}$ | 10,617 | 8,295 | 10,673 | 8,453 | 7,873 |
| $N_{Sum\_split}$ | 7,317 | 11,132 | 8,838 | 11,668 | 6,223 |
| $N_{Bor}$ | 8,805$^{exp}$ | 9,876$^{exp}$ | 10,673 | 8,453 | 8,313$^{exp}$ |
| $N_{Sum}$ | 37,661$^{exp}$ | 43,743$^{exp}$ | 50,401$^{exp}$ | 49,595$^{exp}$ | 54,528$^{exp}$ |
| $T_{split}$ (gen.) | 20,157 | 12,700 | 12,037 | 11,668 | 12,365 |

$N_{anc}$: size of ancestral population; $N_{Bor\_split}$: size of *Pongo pygmaeus* at split; $N_{Sum\_split}$: size of *Pongo abelii* at split; $N_{Bor}$: modern size of *Pongo pygmaeus*; $N_{Sum}$: modern size of *Pongo abelii*; $T_{split}$: time of divergence.
$^{exp}$Exponential growth.

Table S17: The demographic parameters of orangutan history for several models without migrations inferred using *moments* engine in GADMA2. Models differ by set of dynamics used for inference and by the restriction on population sizes. Model follows the restriction (marked by +) when the sum of Bornean and Sumatran population sizes are obliged to equal the ancestral population size before the split. Two histories are also presented in another tables that are indicated on the last row. Ground truth are the simulated parameter values that were obtained from the original paper Locke et al. (2011).

| | Ground truth | ORAN-NOMIG | | ORAN-STRUCT-NOMIG | | |
|---|---|---|---|---|---|---|
| Restriction on population sizes: | | + | − | + | − | − |
| Included dynamics: | | Fixed | Fixed | $^{const}, ^{lin}, ^{exp}$ | $^{const}, ^{lin}, ^{exp}$ | $^{const}, ^{exp}$ |
| Log-likelihood: | | −175,106.12 | −169,881.74 | −174,279.46 | −169,381.38 | −169,956.64 |
| Parameters: | | | | | | |
| $N_{anc}$ | 17,934 | 19,233 | 19,776 | 19,512 | 19,880 | 19,862 |
| $N_{Bor\_split}$ | 10,617 | 8,218 | 6,278 | 10,673 | 6,033 | 6,043 |
| $N_{Sum\_split}$ | 7,317 | 11,014 | 7,753 | 8,838 | 4,515 | 7,692 |
| $N_{Bor}$ | $8{,}805^{exp}$ | 9,455 | $10{,}886^{exp}$ | 10,673 | $10{,}915^{exp}$ | $11{,}012^{exp}$ |
| $N_{Sum}$ | $37{,}661^{exp}$ | $40{,}661^{exp}$ | $56{,}415^{exp}$ | $50{,}401^{exp}$ | $45{,}205^{lin}$ | $57{,}081^{exp}$ |
| $T_{split}$ (gen.) | 20,157 | 12,301 | 11,458 | 12,037 | 11,287 | 11,353 |
| Parameters are also presented in: | | — | Table S14 | Table S16 | Table S18 | — |

$N_{anc}$: size of ancestral population; $N_{Bor\_split}$: size of *Pongo pygmaeus* at split; $N_{Sum\_split}$: size of *Pongo abelii* at split; $N_{Bor}$: modern size of *Pongo pygmaeus*; $N_{Sum}$: modern size of *Pongo abelii*; $T_{split}$: time of divergence.
$^{const}$Constant size.
$^{lin}$Linear growth.
$^{exp}$Exponential growth.

Table S18: The demographic parameters of orangutan history for models with linear size change of Sumatran population and without migrations inferred with different engines in GADMA2. Ground truth are the simulated parameter values that were obtained from the original paper Locke et al. (2011). *Momi2* engine was excluded as it does not support linear size change. Log-likelihood values are not comparable between different engines.

| | Ground truth | ∂a∂i | *moments* | *momentsLD* |
|---|---|---|---|---|
| Log-likelihood: | | −172,654.12 | $-169,381.38$ | −19,321.56 |
| Parameters: | | | | |
| $N_{anc}$ | 17,934 | 19,912 | 19,880 | 19,880 |
| $N_{Bor\_split}$ | 10,617 | 6,559 | 6,033 | 6,056 |
| $N_{Sum\_split}$ | 7,317 | 5,119 | 4,515 | 4,544 |
| $N_{Bor}$ | 8,805 | 11,071 | 10,915 | 11,080 |
| $N_{Sum}$ | $37{,}661^{exp}$ | $46{,}004^{lin}$ | $45{,}205^{lin}$ | $45{,}399^{lin}$ |
| $T_{split}$ (gen.) | 20,157 | 11,937 | 11,287 | 11,384 |

$N_{anc}$: size of ancestral population; $N_{Bor\_split}$: size of *Pongo pygmaeus* at split; $N_{Sum\_split}$: size of *Pongo abelii* at split; $N_{Bor}$: size of *Pongo pygmaeus* after exponential growth; $N_{Sum}$: size of *Pongo abelii*; $T_{split}$: time of divergence.
$^{lin}$Linear growth.
$^{exp}$Exponential growth.

Table S19: Comparison of average times for log-likelihood evaluation between used GADMA2's likelihood engines and models of orangutan history. For each model and likelihood engine mean value and standard deviation are reported.

| Model | ∂a∂i | *moments* | *momi2* | *momentsLD* |
|---|---|---|---|---|
| ORAN-NOMIG | $0.05 \pm 0.14$ | $0.03 \pm 0.01$ | $0.01 \pm 0.006$ | $15.02 \pm 25.71$ |
| ORAN-MIG | $0.24 \pm 2.02$ | $0.16 \pm 0.11$ | – | $27.98 \pm 44.59$ |
| ORAN-STRUCT-NOMIG | $0.07 \pm 0.34$ | $0.06 \pm 0.02$ | $0.02 \pm 0.01$ | $15.73 \pm 27.55$ |
| ORAN-STRUCT-MIG | $0.17 \pm 0.75$ | $0.27 \pm 0.16$ | – | $18.09 \pm 39.26$ |
| ORAN-PULSE1 | – | – | $0.39 \pm 0.16$ | – |
| ORAN-PULSE3 | – | – | $1.18 \pm 0.51$ | – |
| ORAN-PULSE7 | – | – | $3.13 \pm 1.34$ | – |

Table S20: Mean number of log-likelihood evaluations required for demographic inference averaged over eight GADMA2 runs for each engine and model of orangutan history.

| Model | ∂a∂i | *moments* | *momi2* | *momentsLD* |
|---|---|---|---|---|
| ORAN-NOMIG | 11,442 | 8,428 | 6,993 | 10,318 |
| ORAN-MIG | 11,411 | 12,320 | – | 13,570 |
| ORAN-STRUCT-NOMIG | 3,977 | 3,881 | 7,020 | 2,570 |
| ORAN-STRUCT-MIG | 5,356 | 7,200 | – | 10,318 |
| ORAN-PULSE1 | – | – | 7,667 | – |
| ORAN-PULSE3 | – | – | 9,160 | – |
| ORAN-PULSE7 | – | – | 9,178 | – |

### S3 Inference of inbreeding coefficients

#### S3.1 American puma dataset

Table S21: Result statistics obtained from 100 repeats of two ∂a∂i's optimization techniques and GADMA2 in a case of the demographic inference without inbreeding for the American Puma populations. The reported statistics include the mean and standard deviation of the number of evaluations, CPU times, and log-likelihoods. GADMA2 is compared with two ∂a∂i's optimization techniques: single optimization and optimization with multiple restarts. In order to match the number of evaluations with GADMA2, the ∂a∂i optimization with multiple restarts has 18 restarts. Additionally, results from Blischak et al. (2020) are included, which were obtained using single ∂a∂i optimization with different initialization process. BFGS optimization was used as an optimization from ∂a∂i. The results obtained from GADMA2 are highlighted in bold, as they achieved the best mean and best log-likelihood values.

| | Results from Blischak et al. (2020) | Optimization from ∂a∂i 1 restart | 18 restarts | GADMA2 |
|---|---|---|---|---|
| Mean eval. number | Unknown | $238 \pm 53$ | $4{,}307 \pm 200$ | $4{,}103 \pm 1{,}930$ |
| Mean CPU time (min) | Unknown | $0.8 \pm 3.6$ | $17 \pm 18$ | $88 \pm 43$ |
| Best log-likelihood | −453,003.04 | −452,987.45 | −452,987.45 | **−452,969.48** |
| Mean log-likelihood | −585,294 | −1,418,712 | −455,783 | **−453,000** |
| Std log-likelihood | 183,002 | 2,213,096 | 22,275 | 53 |

Table S22: Result statistics obtained from 100 repeats of two ∂a∂i's optimization techniques and GADMA2 in a case of the demographic inference with inbreeding for the American Puma populations. The reported statistics include the mean and standard deviation of the number of evaluations, CPU times, and log-likelihoods. GADMA2 is compared with two ∂a∂i's optimization techniques: single optimization and optimization with multiple restarts. In order to match the number of evaluations with GADMA2, the ∂a∂i optimization with multiple restarts has 16 restarts. Additionally, results from Blischak et al. (2020) are included, which were obtained using single ∂a∂i optimization with different initialization process. BFGS optimization was used as an optimization from ∂a∂i. The results obtained from GADMA2 are highlighted in bold, as they achieved the best mean and best log-likelihood values.

| | Results from Blischak et al. (2020) | Optimization from ∂a∂i | | GADMA2 |
|---|---|---|---|---|
| | | 1 restart | 16 restarts | |
| Mean eval. number | Unknown | $394 \pm 82$ | $6{,}245 \pm 324$ | $6{,}193 \pm 2{,}680$ |
| Mean CPU time (min) | Unknown | $1.3 \pm 1.4$ | $25 \pm 19$ | $93 \pm 47$ |
| Best log-likelihood | −318,058.07 | −317,370.88 | −317,370.88 | **−317,239.49** |
| Mean log-likelihood | −391,303 | −1,729,870 | −320,947 | **−319,451** |
| Std log-likelihood | 121,447 | 4,339,276 | 5,029 | 7,340 |

Table S23: Log-scale standard deviations for parameters in the model without inbreeding for American pumas across a series of step sizes.

| $\epsilon$ | $\sigma_{\log \nu_{TX}}$ | $\sigma_{\log \nu_{FL}}$ | $\sigma_{\log \tau_1}$ | $\sigma_{\log \tau_2}$ | $\sigma_{\log \theta}$ |
|---|---|---|---|---|---|
| $10^{-2}$ | 1.0578 | 1.2988 | 1.5868 | 1.2969 | 0.0484 |
| $10^{-3}$ | 0.0324 | 0.0090 | 0.0763 | 0.0119 | 0.0118 |
| $10^{-4}$ | 0.0917 | 0.0899 | 0.1537 | 0.0920 | 0.0120 |
| $10^{-5}$ | 0.0135 | 0.0116 | 0.0194 | 0.0112 | 0.0104 |
| $10^{-6}$ | 0.0057 | 0.0133 | 0.0015 | 0.0075 | 0.0099 |
| $10^{-7}$ | 1.16e-04 | 7.34e-05 | 3.72e-05 | 7.29e-05 | 9.75e-03 |

Table S24: Log-scale standard deviations for parameters in the model with inbreeding for American pumas across a series of step sizes.

| $\epsilon$ | $\sigma_{\log \nu_{TX}}$ | $\sigma_{\log \nu_{FL}}$ | $\sigma_{\log \tau_1}$ | $\sigma_{\log \tau_2}$ | $\sigma_{\log F_{TX}}$ | $\sigma_{\log F_{FL}}$ | $\sigma_{\log \theta}$ |
|---|---|---|---|---|---|---|---|
| $10^{-2}$ | 0.0798 | 6.0780 | 0.2780 | 5.9785 | 0.0609 | 0.0617 | 0.0394 |
| $10^{-3}$ | 0.0276 | 0.0050 | 0.0391 | 0.0034 | 0.0100 | 0.0092 | 0.0425 |
| $10^{-4}$ | 0.0541 | 0.1511 | 0.0295 | 0.1481 | 0.0065 | 0.0093 | 0.0557 |
| $10^{-5}$ | 0.3870 | 0.2685 | 1.4893 | 0.2638 | 0.0066 | 0.0093 | 0.4978 |
| $10^{-6}$ | 0.0406 | 0.0253 | 0.0192 | 0.0242 | 0.0067 | 0.0093 | 0.0504 |
| $10^{-7}$ | 0.0667 | 0.0078 | 0.0622 | 0.0171 | 0.0476 | 0.0169 | 0.1126 |

Table S25: Maximum likelihood parameters inferred from the demographic models for the Texas and Florida populations of American puma.

| | Model 1 Blischak et al. (2020) | Model 1 GADMA2 | Model 2 Blischak et al. (2020) | Model 2 GADMA2 |
|---|---|---|---|---|
| Number of parameters | 4 | 4 | 6 | 6 |
| Log-likelihood | −453,003.05 | −452,492.70 | −318,058.08 | **−316,115.56** |
| Population size (95% CI) | | | | |
| $N_A$ | 120,000 (92,400 – 157,000) | 118,173 (107,476 – 129,935) | 130,000 (129,000 – 132,000) | 133,934 (123,961 – 144,709) |
| $N_{TX}$ | 23,700 (3,490 – 161,000) | 16,777 (1,930 –145,786) | 70,800 (63,300 – 79,200) | 34,838 (27,594 – 43,983) |
| $N_{FL}$ | 1,210 (118 – 12,500) | 860 (61 – 1,1982) | 1,600 (128 – 19,100) | 374 (0 – 60,398,210) |
| Time in years (95% CI) | | | | |
| $T_1$ | 26,800 (504 – 1,420,000) | 14,833 (604 – 363,950) | 247,000 (169,000 – 359,000) | 387,717 (208,231 – 721,912) |
| $T_2$ | 8,230 (784 – 86,500) | 5,806 (418 – 80,583) | 7,820 (650 – 94,200) | 1,836 (0 – 243,391,796) |
| Inbreeding coefficients (95% CI) | | | | |
| $F_{TX}$ | NA | NA | 0.440 (0.408 – 0.474) | 0.454 (0.403 – 0.512) |
| $F_{FL}$ | NA | NA | 0.607 (0.588 – 0.626) | 0.627 (0.556 – 0.708) |

$N_A$: size of ancestral population; $N_{TX}$: size of ancestral population after growth and size of Texas population; $N_{FL}$: size of Florida population after divergence; $T_1$: time of epoch between ancestral population size growth and split event; $T_2$: time of divergence; $F_{TX}$: inbreeding coefficient for Texas population; $F_{FL}$: inbreeding coefficient for Florida population.

#### S3.2 Domesticated cabbage dataset

Table S26: Result statistics obtained from 100 repeats of two ∂a∂i's optimization techniques and GADMA2 in a case of the demographic inference without inbreeding for the domesticated cabbage. The reported statistics include the mean and standard deviation of the number of evaluations, CPU times, and log-likelihoods. GADMA2 is compared with two ∂a∂i's optimization techniques: single optimization and optimization with multiple restarts. Additionally, results from Blischak et al. (2020) are included, which were obtained using single ∂a∂i optimization with different initialization process. In order to match the number of evaluations with GADMA2, the ∂a∂i optimization with multiple restarts has 27 restarts. BOBYQA optimization was used as an optimization from ∂a∂i. The results obtained from ∂a∂i's optimization with multiple restarts are highlighted in bold, as they achieved the best mean and best log-likelihood values.

| | Results from Blischak et al. (2020) | Optimization from ∂a∂i | | GADMA2 |
|---|---|---|---|---|
| | | 1 restart | 27 restarts | |
| Mean eval. number | Unknown | $205 \pm 307$ | $6{,}053 \pm 1,733$ | $6{,}034 \pm 2,764$ |
| Mean CPU time (min) | Unknown | $6 \pm 19$ | $160 \pm 131$ | $84 \pm 40$ |
| Best log-likelihood | −24,330.40 | −24,303.37 | **−24,292.72** | −24,307.64 |
| Mean log-likelihood | −45,431 | −55,567 | **−25,384** | −25,723 |
| Std log-likelihood | 12,678 | 19,498 | 4,371 | 2,689 |

Table S27: Result statistics obtained from 100 repeats of two ∂a∂i's optimization techniques and GADMA2 in a case of the demographic inference with inbreeding for the domesticated cabbage. The reported statistics include the mean and standard deviation of the number of evaluations, CPU times, and log-likelihoods. GADMA2 is compared with two ∂a∂i's optimization techniques: single optimization and optimization with multiple restarts. Additionally, results from Blischak et al. (2020) are included, which were obtained using single ∂a∂i optimization with different initialization process. In order to match the number of evaluations with GADMA2, the ∂a∂i optimization with multiple restarts has 16 restarts. BOBYQA optimization was used as an optimization from ∂a∂i. The results obtained from ∂a∂i's optimization with multiple restarts are highlighted in bold, as they achieved the best mean and slightly better than GADMA2 best log-likelihood values.

| | Results from Blischak et al. (2020) | Optimization from ∂a∂i | | GADMA2 |
|---|---|---|---|---|
| | | 1 restart | 12 restarts | |
| Mean eval. number | Unknown | $534 \pm 773$ | $5{,}923 \pm 2{,}501$ | $5{,}872 \pm 3{,}282$ |
| Mean CPU time (min) | Unknown | $33 \pm 46$ | $357 \pm 173$ | $418 \pm 232$ |
| Best log-likelihood | −4,281.14 | −4,271.25 | **−4,270.35** | −4,270.37 |
| Mean log-likelihood | −10,159 | −26,023 | **−4,398** | −4,677 |
| Std log-likelihood | 48,884 | 29,721 | 285 | 778 |

Table S28: Log-scale standard deviations for parameters in the model without inbreeding for domesticated cabbage across a series of step sizes.

| $\epsilon$ | $\sigma_{\log \nu_1}$ | $\sigma_{\log \nu_2}$ | $\sigma_{\log \tau_1}$ | $\sigma_{\log \tau_2}$ | $\sigma_{\log \theta}$ |
|---|---|---|---|---|---|
| $10^{-2}$ | 3.8339 | 7.1222 | 0.9787 | 9.4997 | 0.2319 |
| $10^{-3}$ | 4.5392 | 15.1865 | 1.5670 | 13.8694 | 0.1161 |
| $10^{-4}$ | 10.6951 | 38.0201 | 3.1475 | 41.1826 | 0.1824 |
| $10^{-5}$ | 3.3863 | 7.3402 | 1.0582 | 6.2260 | 0.0642 |
| $10^{-6}$ | 0.0989 | 0.1912 | 0.0926 | 0.1869 | 0.0195 |
| $10^{-7}$ | 0.0095 | 0.0317 | 0.0651 | 0.0605 | 0.0302 |

Table S29: Log-scale standard deviations for parameters in the model with inbreeding for domesticated cabbage across a series of step sizes.

| $\epsilon$ | $\sigma_{\log \nu_1}$ | $\sigma_{\log \nu_2}$ | $\sigma_{\log \tau_1}$ | $\sigma_{\log \tau_2}$ | $\sigma_{\log F}$ | $\sigma_{\log \theta}$ |
|---|---|---|---|---|---|---|
| $10^{-2}$ | 0.0412 | 1.5233 | 0.2200 | 0.3345 | 0.0190 | 0.0319 |
| $10^{-3}$ | 0.0412 | 0.9052 | 0.3668 | 0.3451 | 0.0186 | 0.0592 |
| $10^{-4}$ | 0.0412 | 0.8094 | 0.2834 | 0.2754 | 0.0188 | 0.0460 |
| $10^{-5}$ | 0.0412 | 1.2076 | 0.2753 | 0.2676 | 0.0188 | 0.0448 |
| $10^{-6}$ | 0.0430 | 1.6379 | 0.2079 | 0.1165 | 0.0195 | 0.0347 |
| $10^{-7}$ | 1.4932 | 1.4230 | 1.8616 | 0.3920 | 0.1200 | 0.5023 |

Table S30: Maximum likelihood parameters inferred from the demographic models for the domesticated cabbage population.

| | Model 1 Blischak et al. (2020) | Model 1 GADMA2 | Model 2 Blischak et al. (2020) | Model 2 GADMA2 |
|---|---|---|---|---|
| Number of parameters | 5 | 5 | 6 | 6 |
| Log-likelihood | −24,330.40 | −24,137.34 | −4,281.14 | **−4,267.32** |
| Population size (95% CI) | | | | |
| $N_A$ | 19,100 (18,500 – 19,800) | 19,121 (12,136 – 30,128) | 17,500 (16,900 – 18,100) | 17,496 (16,432 – 18,628) |
| $N_1$ | 123,000 (80,400 – 190,000) | 95,047 (81 – 111,047,156) | 31,600 (28,900 – 34,700) | 31,792 (28,700 – 35,218) |
| $N_2$ | 592 (547 – 641) | 10 (0 – 7,848,832) | 215,000 (4,910 – 9,370,000) | 174,961,828 (8,384,199 – 3,651,110,690) |
| Time in years (95% CI) | | | | |
| $T_1$ | 5,870 (5,200 – 6,620) | 6,128 (1,416 – 26,525) | 16,600 (12,900 – 21,200) | 16,552 (11,330 – 24,180) |
| $T_2$ | 38.3 (32.5 – 45.1) | 0.616 (0 – 47,943,409) | 322 (94.2 – 1,097) | 256 (139 – 471) |
| Inbreeding coefficient (95% CI) | | | | |
| $F$ | NA | NA | 0.578 (0.557 – 0.599) | 0.577 (0.556 – 0.599) |

$N_A$: size of ancestral population; $N_1$: size of population during the first epoch; $N_2$: size of population during the first epoch; $T_1$: time of the first epoch; $T_2$: time of the second epoch; $F$: inbreeding coefficient.
